# Structural basis for the specificity of PPM1H phosphatase for Rab GTPases

**DOI:** 10.1101/2021.02.17.431620

**Authors:** Dieter Waschbüsch, Kerryn Berndsen, Pawel Lis, Axel Knebel, Yuko P. Y. Lam, Dario R. Alessi, Amir R. Khan

## Abstract

LRRK2 serine/threonine kinase is associated with inherited Parkinson’s disease. LRRK2 phosphorylates a subset of Rab GTPases within their switch 2 motif to control their interactions with effectors. Recent work has shown that the metal-dependent protein phosphatase PPM1H counteracts LRRK2 by dephosphorylating Rabs. PPM1H is highly selective for LRRK2 phosphorylated Rabs, and closely related PPM1J exhibits no activity toward substrates such as Rab8a phosphorylated at Thr72 (pThr72). Here we have identified the structural determinant of PPM1H specificity for Rabs. The crystal structure of PPM1H reveals a structurally conserved phosphatase fold that strikingly has evolved a 110-residue flap domain adjacent to the active site. The flap domain distantly resembles tudor domains that interact with histones in the context of epigenetics. Cellular assays, crosslinking and 3-D modelling suggest that the flap domain encodes the docking motif for phosphorylated Rabs. Consistent with this hypothesis, a PPM1J chimera with the PPM1H flap domain dephosphorylates pThr72 of Rab8a both *in vitro* and in cellular assays. Therefore, PPM1H has acquired a Rab-specific interaction domain within a conserved phosphatase fold.

## Introduction

Metal-dependent Ser/Thr phosphatases (PPMs) have a structurally conserved catalytic domain that adopts a β-sandwich fold with Mg^2+^/Mn^2+^ ions at the active site. Among the family of human enzymes is PPM1A (formerly PP2Cα), which reverses stress-mediated protein kinase cascades [1–3]; PHLPP1/2 which regulates AGC kinases and cellular homeostasis[4]; and pyruvate dehydrogenase phosphatase (PDP1) that is expressed in the mitochondrial matrix and regulates the activity of pyruvate dehydrogenase in metabolism[5, 6]. The core PPM fold, first identified by the structure of PPM1A[7], consists of an 11-stranded β-sandwich flanked on both sides by α-helices. The catalytic cleft is formed on one side of the β-sandwich and comprises a binuclear Mg^2+^/Mn^2+^ metal center that is coordinated by conserved aspartate residues. The 250-residue PPM fold is better conserved in structure rather than sequence (20-50% identities) across the mammalian enzymes. There is also considerable diversity among PPMs involving the incorporation of sequence elements outside of the catalytic domain. For example, PPM1A has a C-terminal α-helical domain that is not required for catalysis but may contribute to substrate specificity[7–9]. Several enzymes including PPM1A, PPM1B, PPM1K and PDP1 also have a short 50-residue insertion termed the ‘flap’ subdomain that is poorly conserved in sequence and structure. This region is predicted to contribute to substrate specificity, although chimeric enzymes involving grafts of the flap have not been successful in transferring substrate preference[10]. Studies of bacterial enzymes have proposed a third metal-binding site that contributes to catalysis *via* coordination with a conserved aspartate residue [11–13]. Mutation of the equivalent residue in human PPM1A to alanine (D146A) enabled trapping of a complex of PPM1A with a cyclic phospho-peptide and subsequent structure determination[9].

Recently, PPM1H phosphatase has been identified as the enzyme that counteracts the LRRK2 signaling cascade by dephosphorylating Rab GTPases[14]. A subset of at least 7 Rabs are physiological substrates of LRRK2[15, 16], a Ser/Thr kinase that is associated with inherited and sporadic forms of Parkinson’s disease (PD)[17, 18]. Rabs are members of the Ras superfamily of molecular switches that regulate membrane trafficking in eukaryotes. Rabs oscillate between a membrane-bound GTP form and cytosolic GDP form that is distinguished by local conformational changes in nucleotide-sensitive switch 1 and switch 2 motifs[19]. LRRK2 phosphorylates Rab8a and Rab10 at a conserved threonine residue in their switch 2 α-helix (pThr72 in Rab8a and pThr73 in Rab10). Phosphorylated Rab8a/10 (pRab8a/10) recruit phospho-specific effectors RILPL1 and RILPL2 (Rab interacting lysosomal protein-like 1 and 2) to subcellular compartments, downstream of LRRK2 activation. Autosomal dominant PD mutations that activate LRRK2 kinase interfere with the formation of primary cilia through a pathway involving pRab8a/10 binding to RILPL1[20]. LRRK2 kinase inhibitors are currently in phase 1 and 2 clinical trials with PD patients[21], while alternative strategies to antagonize LRRK2 signaling could be beneficial for future therapeutics.

Here we describe the crystal structure of PPM1H, the phosphatase that counteracts the LRRK2 pathway. The structure reveals novel features that have been incorporated into the core catalytic domain. The first is a 110-residue ‘flap domain’ that is positioned next to the catalytic cleft. This domain is an expansion of a 50-residue flap that is found in other members of the PPM family. The PPM1H flap domain adopts an α/β fold resembling tudor domains that regulate histone functions in an epigenetic context. On the opposite face of the cleft, a 3-stranded β-sheet motif (β-motif) is also inserted into the core PPM fold. Thirdly, PPM1H has an N-terminal extension that winds behind the active site and inserts into the hydrophobic core of the β-sandwich. This anchor-like interaction has apparently evolved in the PPM1H/J/M subfamily of phosphatases as a motif that contributes to folding of the enzymes. Mutagenesis, cellular assays, crosslinking and modelling studies of a phospho-Rab substrate into the active site suggest that the PPM1H flap domain forms a docking site for phosphorylated Rab GTPases. In support of this hypothesis, transfer of the flap domain of PPM1H onto PPM1J is sufficient to convert the PPM1J chimera into an active phospho-Rab8a phosphatase. Therefore, PPM1H phosphatase has evolved substrate specificity through the acquisition of a Rab-specific flap domain within the framework of a conserved catalytic domain.

## Results and Discussion

### Overall structure of PPM1H phosphatase

Full-length PPM1H expressed in *E.coli* cells was unstable and prone to degradation, thus difficult to crystallize. To design a crystallisable protein, the N-terminal residues 1-32 were eliminated due to predicted flexibility (Fig 1A). We also introduced a D288A mutation (PPM1H^DA^) that destabilizes a third metal ion (Mg^2+^/Mn^2+^) at the active site [9], which previous studies showed acts as a substrate trapping mutant. Crystals were grown of PPM1H^DA^ that diffracted to 3.1 Å resolution (Table 1), but no crystals grew of the WT enzyme. To grow WT crystals and improve diffraction, a ‘loop deletion’ variant of PPM1H (PPM1H-LD) was designed to eliminate a flexible and non-conserved loop (188-226) that was predicted to be distant from the active site. WT and D288A variants of PPM1H (PPM1H^WT^-LD, PPM1H^DA^-LD) diffracted to 2.5 Å and 2.6 Å resolution, respectively. The variants PPM1H^DA^, PPM1H^WT^-LD, and PPM1H^DA^-LD have identical 3-D structures with two Mg^2+^ ions at the active site. In addition to the Mg^2+^ complexes, a structure of PPM1H^WT^ (33-514, loopDEL) with 3 Mn^2+^ ions was also determined at 2.2 Å resolution (MnPPM1H^WT^-LD). The 2.5 Å model of PPM1H^WT^-LD will be used for ensuing discussions (Fig 1), except for 3-D docking analyses with model substrates in which we used the MnPPM1H^WT^-LD variant. However, all of the structures are identical with only minor differences arising from flexible loops and the presence or absence of the third metal ion at the active site. Deletion of the loop 188-226 and the N-terminus (1-32) does not affect the ability of PPM1H^WT^-LD to dephosphorylate pRab8a relative to the full-length enzyme *in vitro* and in cells (Fig EV1A). Statistics of data collection and refinement for all structures are shown in Table 1.

**Figure 1:**
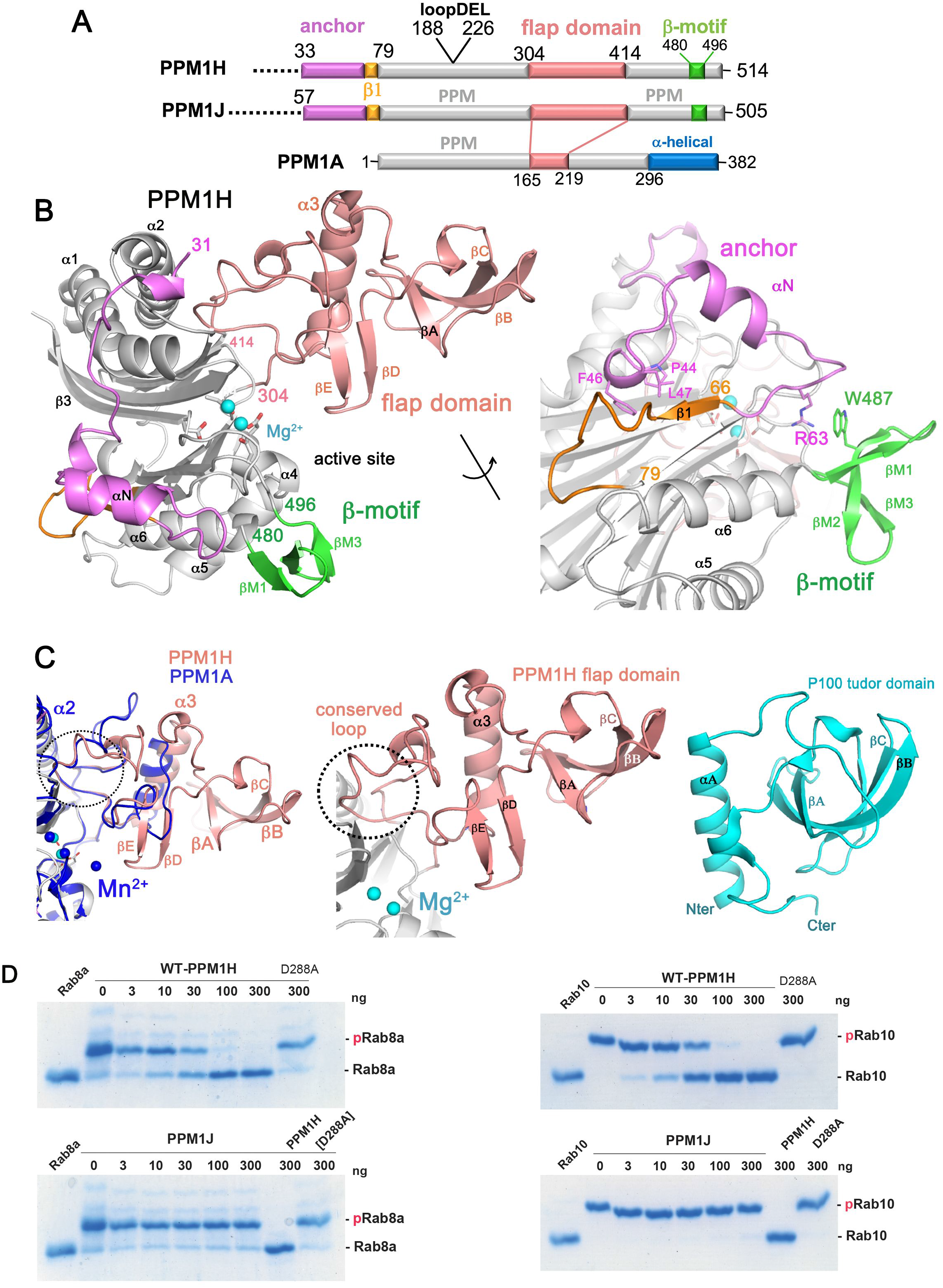
Structure of Rab-specific PPM1H phosphatase. **(A)** Domain organization of PPM1H, PPM1J, and PPM1A. The annotation of regions (anchor, flap domain) is discussed in the text. The loop deletion (188-226) that connects that was engineered to improve diffraction is indicated. **(B)** Ribbon model of the enzyme with a view to the catalytic cleft that contains two Mg^2+^ ions (cyan spheres). The N-terminal region is magenta (33-71), the flap domain is a wheat colour, and the β-sheet motif is green. The loop deletion (188-226) connects α1/α2 on the opposite face relative to the active site. The back view of the enzyme is also shown with a 180° rotation around the axis indicated. Parts of the anchor (RPxFL motif, magenta) that interact with the globular core are shown as stick models, and discussed in the text. The β1 strand is orange to emphasize its non-canonical conformation due to the presence of the preceding anchor. **(C)** Comparisons of the flap domain of PPM1H with PPM1A (left) and the tudor domain (right). Apart from a conserved loop (dotted circle) which forms an interface with the catalytic domain, the sequences and structures of flaps are diverse among the PPM family. **(D)** The indicated amounts of recombinant wild-type and mutant PPM1H or PPM1J (with a His-Sumo N-terminal tag, expressed in *E. coli*) were incubated *in vitro* with 2.5 µg pThr72 phosphorylated Rab8a (*left*) or pThr73 phosphorylated Rab10 (*right*) for 20 min in the presence of 10 mM MgCl_2_ in 40 mM HEPES pH 7.5 buffer. Reactions were terminated by addition of SDS Sample Buffer and analyzed by Phos-tag gel electrophoresis that separates phosphorylated (slow migrating) and dephosphorylated Rabs. The gel was stained with Instant Blue Coomassie. D288A is a substrate-trapping (inactive) variant of PPM1H and was used as a control.

**Table 1.**
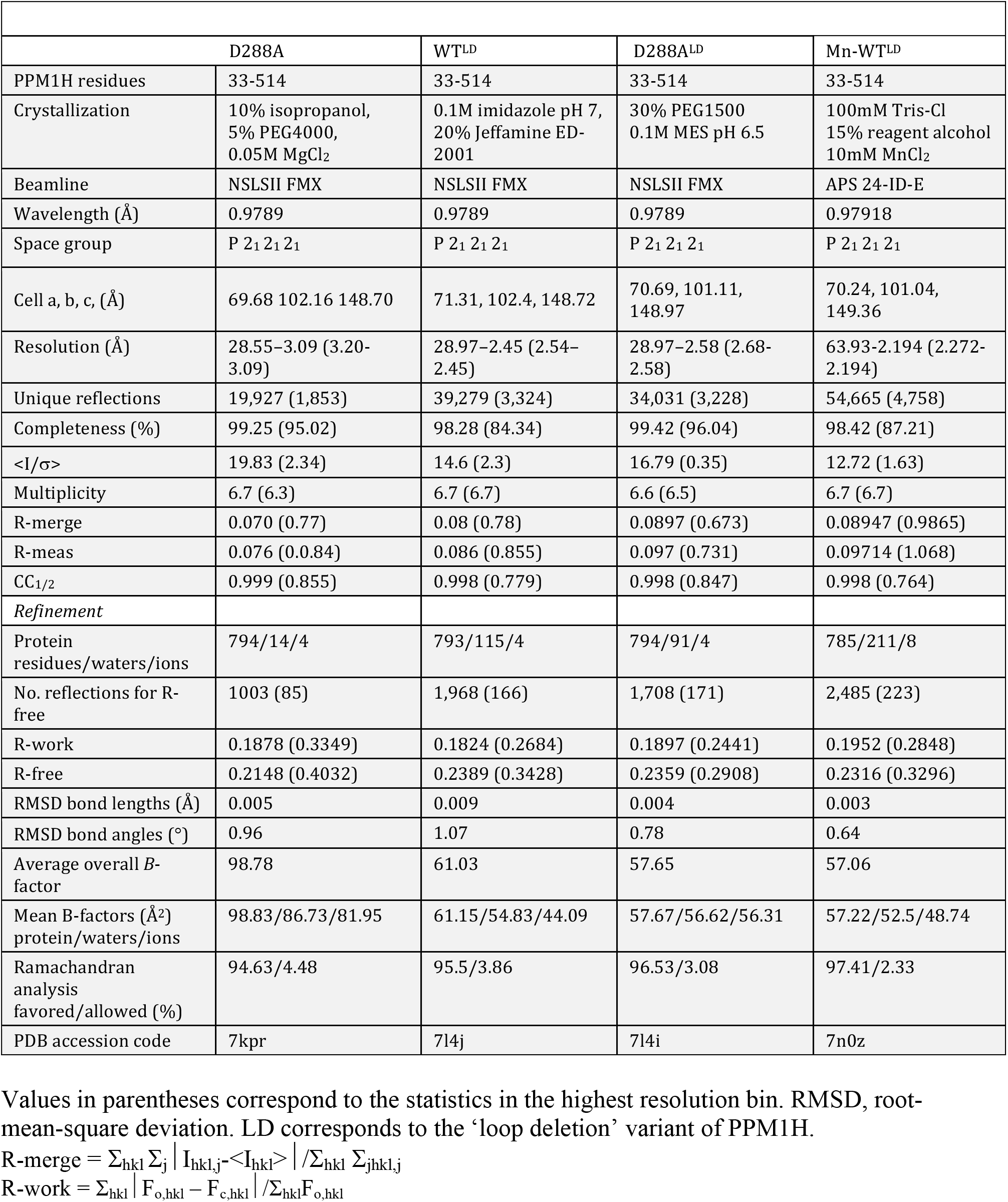
PPM1H Crystallographic Data and Refinement Statistics

| Table 1. PPM1H Crystallographic Data and Refinement Statistics |  |  |  |  |
| --- | --- | --- | --- | --- |
|  | D288A | WT<sup>LD</sup> | D288A<sup>LD</sup> | Mn-WT<sup>LD</sup> |
| PPM1H residues | 33-514 | 33-514 | 33-514 | 33-514 |
| Crystallization | 10% isopropanol, 5% PEG4000, 0.05M MgCl<sub>2</sub> | 0.1M imidazole pH 7, 20% Jeffamine ED-2001 | 30% PEG1500 0.1M MES pH 6.5 | 100mM Tris-Cl 15% reagent alcohol 10mM MnCl<sub>2</sub> |
| Beamline | NSLSII FMX | NSLSII FMX | NSLSII FMX | APS 24-ID-E |
| Wavelength (Å) | 0.9789 | 0.9789 | 0.9789 | 0.97918 |
| Space group | P 2<sub>1</sub> 2<sub>1</sub> 2<sub>1</sub> | P 2<sub>1</sub> 2<sub>1</sub> 2<sub>1</sub> | P 2<sub>1</sub> 2<sub>1</sub> 2<sub>1</sub> | P 2<sub>1</sub> 2<sub>1</sub> 2<sub>1</sub> |
| Cell a, b, c, (Å) | 69.68 102.16 148.70 | 71.31, 102.4, 148.72 | 70.69, 101.11, 148.97 | 70.24, 101.04, 149.36 |
| Resolution (Å) | 28.55–3.09 (3.20–3.09) | 28.97–2.45 (2.54–2.45) | 28.97–2.58 (2.68–2.58) | 63.93–2.194 (2.272–2.194) |
| Unique reflections | 19,927 (1,853) | 39,279 (3,324) | 34,031 (3,228) | 54,665 (4,758) |
| Completeness (%) | 99.25 (95.02) | 98.28 (84.34) | 99.42 (96.04) | 98.42 (87.21) |
| <I/σ> | 19.83 (2.34) | 14.6 (2.3) | 16.79 (0.35) | 12.72 (1.63) |
| Multiplicity | 6.7 (6.3) | 6.7 (6.7) | 6.6 (6.5) | 6.7 (6.7) |
| R-merge | 0.070 (0.77) | 0.08 (0.78) | 0.0897 (0.673) | 0.08947 (0.9865) |
| R-meas | 0.076 (0.84) | 0.086 (0.855) | 0.097 (0.731) | 0.09714 (1.068) |
| CC<sub>1/2</sub> | 0.999 (0.855) | 0.998 (0.779) | 0.998 (0.847) | 0.998 (0.764) |
| *Refinement* |  |  |  |  |
| Protein residues/waters/ions | 794/14/4 | 793/115/4 | 794/91/4 | 785/211/8 |
| No. reflections for R-free | 1003 (85) | 1,968 (166) | 1,708 (171) | 2,485 (223) |
| R-work | 0.1878 (0.3349) | 0.1824 (0.2684) | 0.1897 (0.2441) | 0.1952 (0.2848) |
| R-free | 0.2148 (0.4032) | 0.2389 (0.3428) | 0.2359 (0.2908) | 0.2316 (0.3296) |
| RMSD bond lengths (Å) | 0.005 | 0.009 | 0.004 | 0.003 |
| RMSD bond angles (°) | 0.96 | 1.07 | 0.78 | 0.64 |
| Average overall B-factor | 98.78 | 61.03 | 57.65 | 57.06 |
| Mean B-factors (Å<sup>2</sup>) protein/waters/ions | 98.83/86.73/81.95 | 61.15/54.83/44.09 | 57.67/56.62/56.31 | 57.22/52.5/48.74 |
| Ramachandran analysis favored/allowed (%) | 94.63/4.48 | 95.5/3.86 | 96.53/3.08 | 97.41/2.33 |
| PDB accession code | 7kpr | 7l4j | 7l4i | 7n0z |

Values in parentheses correspond to the statistics in the highest resolution bin. RMSD, root-mean-square deviation. LD corresponds to the ‘loop deletion’ variant of PPM1H.

The structure of PPM1H adopts a conserved PPM fold consisting of 10 β-strands organized into a 5×5 β-sandwich (Fig 1B). The first β-sheet comprises β-strands β2, β7, β8, β10 and β11. The second β-sheet consists of β-strands β3, β4, β5, β6, and β9. Two α-helices pack against the each of the two concave surfaces of the β-sandwich. The long and curved α-helices α1 and α2 are oriented in an anti-parallel manner and pack against the second β-sheet. The shorter helices α4 and α5 are also oriented in an anti-parallel fashion and pack against the first β-sheet. The active site cleft is formed by loops, including a short α-helical loop (α4, residues 443-447), that connect β-strands on one edge of the β-sandwich. Three conserved aspartate residues – D151, D437 and D498 – form direct contacts with two Mg^2+^ ions (M1 and M2) at the active site. M1 is coordinated by D437 and D498, while D151 coordinates both metal sites. The metal binding includes water molecules, and the geometry is approximately octahedral for both metal sites (M1 and M2). A structure-based sequence alignment of PPM family enzymes is shown with secondary structures and other annotations (Fig EV1B).

One of the distinctive features of PPM1H is a 110-residue flap domain that adopts an α/β fold (Fig 1C). All known structures of human PPMs (PPM1A, PPM1B, PPM1K and PDP1) have a shorter 50-residue flap that is inserted between the terminal β-strands of the two β-sheets (β8 and β9 in PPM1H). In PPM1H, this domain comprises an α-helix that stacks against a highly twisted β-sheet. This extended flap domain effectively creates a surface adjacent to the active site for potential substrate recognition. Matching of the flap domain to structures in the DALI server [22] reveals a resemblance to the tudor domain despite the absence of sequence similarities (Fig 1C). Tudor domains are also composed of an α-helix that stacks against a highly twisted anti-parallel β-sheet. These domains are involved in the recognition of methylated lysine or arginine residues of histones *via* an aromatic cage. Although roughly similar in appearance, the α/β flap domain of PPM1H has a different topology and lacks characteristic structural motifs such as the aromatic cage that are common to tudor domains. Despite flap sequence and structural diversity among the PPM family, a short loop 386-396 that packs against the catalytic domain is highly conserved in sequence and conformation (dotted circle, Fig 1C). This conserved loop is investigated in more detail below.

Compared to structures of other known enzymes, PPM1H has two additional structural elements that are novel. The N-terminal residues 33-79 follow an irregular path behind the active site that spans the two β-sheets of the core catalytic domain (Fig 1B). This region is termed an ‘anchor’ due to a short 3_10_ helix (residues 43-47) that caps the hydrophobic core of the β-barrel. The second feature is a β-sheet motif (β-motif; residues 480-496) that consists of 3 short anti-parallel β-strands. These novel features of PPM1H, along with an expanded flap domain, are shared by the PPM1H/J/M subfamily of phosphatases. The specificity of PPM1H against PPM1J was compared over a 20-minute reaction at room temperature using catalytic amounts of enzyme. These *in vitro* assays used phospho-Rab8a (pRab8a) and pRab10 as substrates, and PhosTag gels [23] to assess dephosphorylation of substrates (Fig 1D). Despite sharing common domains, PPM1J displays no activity, while PPM1H completely dephosphorylates pRab8a and pRab10 under these conditions.

### Crosslinking and 3-D docking suggest that the PPM1H flap is a pRab recognition domain

Purified complexes of pRab8a and the substrate-trapping D288A variant of PPM1H were incubated together in the presence of DSBU (disuccinimidyl dibutyric urea), a mass spectrometry cleavable amine reactive crosslinker that is widely used to identify and map sites of protein-protein interactions [24]. This reagent crosslinks Lys residues to acidic and hydroxyl amino acids located within 32 Å [25]. In the presence of DSBU, pRab8a and PPM1H formed a stoichiometrically heavier band on Coomassie stained gels migrating at ∼140 kDa (Fig 2A). The crosslinking was dependent on phosphorylation of Thr72, since the heavier band failed to form with PPM1H/Rab8a or PPM1H alone incubations. The size of the crosslinked species implied 2 molecules of PPM1H (50 kDa each) and 2 molecules of pRab8a (20 kDa each) in solution. Consistent with dimeric complexes of the enzyme, crosslinked PPM1H alone migrated as a dimer (100 kDa) and further information below is supportive of PPM1H being a dimer. Crosslinked samples were digested under 3 conditions (trypsin, trypsin/Asp-N and trypsin/Glu-C). In addition, SCX cartridge purification was applied in one of the tryptic digested samples to further enrich the crosslinked peptides. PPM1H and pRab8a crosslinked peptides were identified using meroX software[26]. Potential crosslinked peptides with score higher than 50, as well as false discovery rate (FDR) less than 5%, were manually inspected to confirm only a single crosslinked site was proposed from each peptide[27]. This protocol identified numerous residues within the flap domain of PPM1H that were crosslinked to pRab8a (Fig 2B **and Table EV2**). Interestingly, other regions in PPM1H that formed significant crosslinks with pRab8a were situated in likely more flexible regions namely the N-terminal non-catalytic region, and within the 104-142 and 183-235 flexible loops (Fig 2B). These data are consistent with the flap domain of PPM1H functioning as a major substrate recognition site for pRab8a.

**Figure 2:**
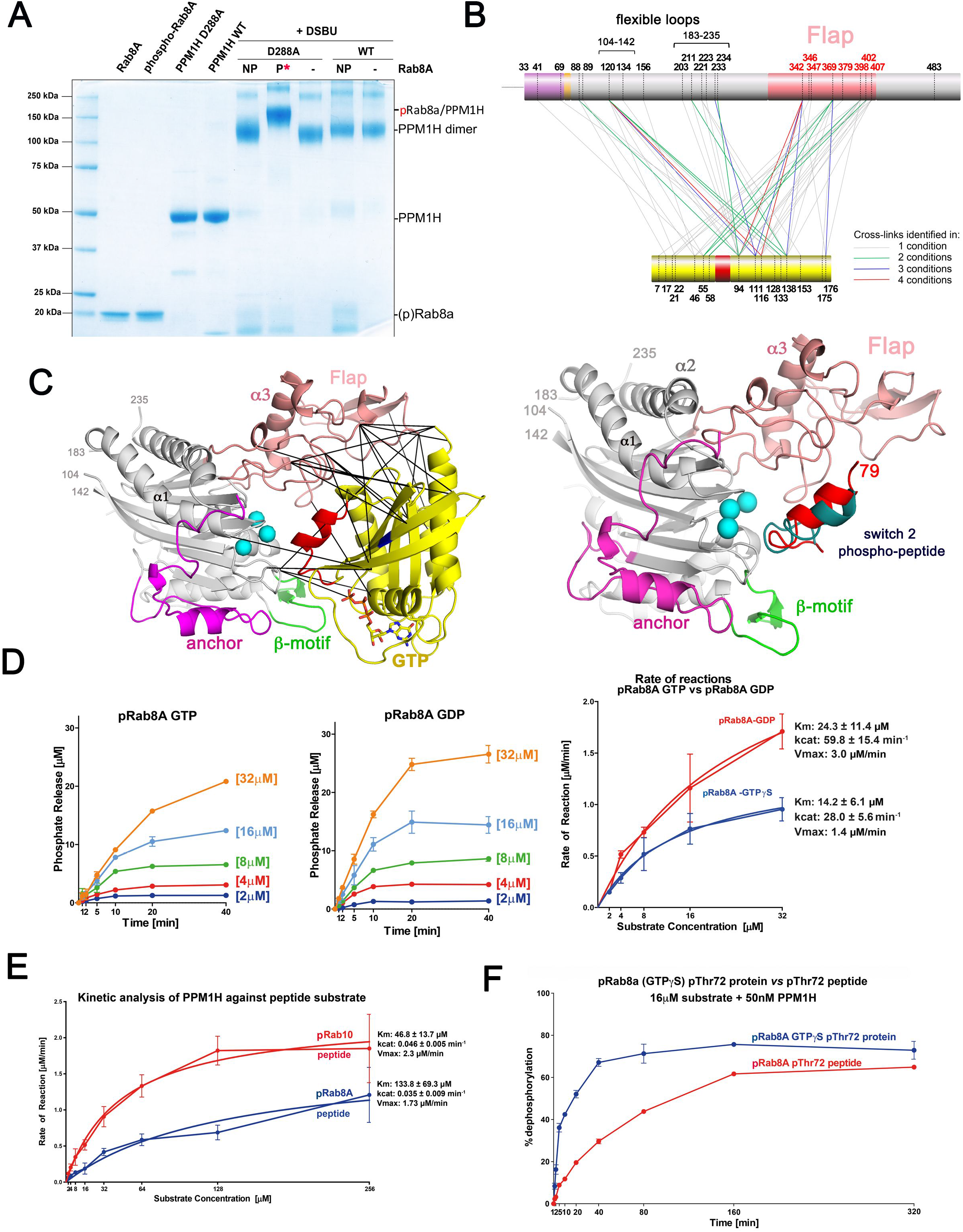
Crosslinking and docking analyses suggest flap domain binds to pRab8a. **(A)** SDS-PAGE page analysis of the PPM1H(D288A): pRab8a complex in the presence of DSBU crosslinker. Control migration of proteins are on the left. NP, non-phosphorylated Rab8a; P, phosphorylated Rab8a. The migration of control and crosslinked proteins is marked on the right. **(B)** Crosslinked peptides from PPM1H and pRab8a are mapped onto the sequence. The flap domain forms extensive crosslinks with pRab8a. In addition, two flexible loops (104-142, 188-226) also have multiple crosslinks with the substrate. **(C)** Ribbon model of pRab8a (*left*) and the switch 2 phosphopeptide (*right*) docked onto the active site of PPM1H. The crosslinks shown between PPM1H and pRab8a are within accepted distance constraints (32 Å) for DSBU [42]. **(D)** Kinetics of phosphate hydrolysis. 25 nM recombinant wild-type PPM1H was incubated with increasing concentrations of pThr72 phosphorylated Rab8a (GTPγS or GDP) as described in Materials and Methods. Initial velocity (V_0_) was calculated by dividing the concentration of released phosphatase (µM) by time (min) and plotted against substrate concentration for pThr72 phosphorylated Rab8a[GTP bound conformation] (*blue*) and pThr72 phosphorylated Rab8a[GDP bound conformation] (*red*). Kinetic constants (K_cat_, V_max_, K_m_) were obtained using GraphPad software. Uncertainties (±) correspond to the standard error of measurements. **(E)** Kinetic analysis of PPM1H as in **(D)** using 50 nM PPM1H and phosphopeptide substrates, as described in Materials and Methods. Initial velocity (V_0_) was calculated by dividing the concentration of phosphatase (µM) by time (min) and plotted against substrate concentration for pThr72 Rab8a phospho-peptide (*blue*) and Rab10 pThr73 phospho-peptide (*red*). **(F)** Side by side comparison of the catalytic activity against protein and peptide by *in vitro* malachite green time course analysis. 50 nM recombinant wild-type PPM1H was incubated with 16 µM Rab8a GTPγS pThr72 phosphorylated protein (*blue*) or Rab8a pThr72 phosphorylated peptide (*red*) for indicated times and analysed as described in Materials and Methods.

In order to further understand the mode of pRab8a interactions with PPM1H, docking analyses were performed. Phosphorylated Rab8a (pRab8a, GTP-bound) from the recent structure of the pRab8a:RILPL2 complex[28] was docked onto PPM1H using Haddock software[29, 30]. The structure of MnPPM1H^WT^-LD with 3 Mn^2+^ ions was used for docking with distance restraints applied between pThr72 of pRab8a and the metal ions of PPM1H. The structures of PPM1H with two Mg^2+^ ions failed to dock with pRab8a at the active site, presumably due to the density of negative charges in the absence of a third metal ion. The top pRab8a docking solution is shown along with the experimentally determined crosslinks from the flap domain (Fig 2C).

The model reveals that the extended active site cleft involving the flap domain forms multiple interactions with pRab8A within the PPM1H active site. Overall, the crosslinking data and docking model are consistent with pRab8a recognition by the flap domain with phosphorylated Thr72 in the switch II motif oriented towards the active site for dephosphorylation.

We also performed docking of the switch 2 α-helical peptide from pRab8a with MnPPM1H^WT^-LD. The 15 residue fragment from 65-79 was extracted from the structure of the pRab8a complex and the N/C termini were capped (acetyl/amide) to eliminate charges. The docking solutions were more heterogenous and many poses would be sterically incompatible with the active site in the context of the full G domain of phospho-Rabs. However, two of the top 10 solutions revealed a similar orientation of the phosphorylated switch 2 helix to the protein/protein dock (Fig 2C). Overall, the complete 8/10 docking solutions of PPM1H/peptide and the top 5/6 docking solutions of PPM1H/protein suggest that the flap domain is very likely to contribute to substrate recognition (Fig EV2A/B).

### PPM1H dephosphorylates pRab8a both in the GDP and GTP states

It is unknown whether PPM1H dephosphorylates pRab8a in its GTP or GDP state. To address this question, we generated pRab8a complexed to GDP or GTPγS and undertook K_m_/V_max_ kinetic analyses (Fig 2D). This revealed that PPM1H dephosphorylated both the GDP and GTP pRab8a with moderately different kinetics. PPM1H dephosphorylated the GDP-pRab8a complex with a K_m_ of 24 μM and a V_max_ of 3.0 μM/min, and GTPγS - pRab8a complex with a K_m_ of 14 μM and a V_max_ of 1.4 μM/min. Therefore, the V_max_ is doubled for the GDP complex, but the GTPγS complex is dephosphorylated with an ∼40% lower K_m_. This suggests that PPM1H *in vivo* would be able to act on both the GTP and GDP states of -pRab proteins. The kinetic parameters for Thr72/Thr73 phosphorylated switch 2 peptides from Rab8a and Rab10 (pRab8a=AGQERF<u>R</u>T*ITT<u>A</u>YYR; pRab10=AGQERF<u>H</u>T*ITT<u>S</u>YYR; residues differing between the two peptides are underlined) that encompass residues 65 - 79 of human Rab8a/10 (Rab8a numbering) were also determined (Fig 2E). The relatively high K_m_ values for peptides (Rab8a=134μM; Rab10=47μM) suggest that they are not optimal substrates compared to the G domain of pRabs. In a side-by-side comparison of substrates assayed at 16 μM, pRab8a (GTPγS) is 7-fold more efficient as a substrate in early time points (5-10 min) relative to the peptide variant (Fig 2F). As a control, there was no significant PPM1H hydrolysis of GTPγS and GDP from Rab complexes under the conditions of these assays (Fig EV2C).

### PPM1H is a dimer

The asymmetric unit in all 4 structures is a dimer that is facilitated by interactions between α3 of the flap domains, as well as contacts between the flap domain (residues 356-360) and α2 of the catalytic domain (Fig 3A). Given the observed dimeric state of PPM1H with DSBU crosslinker, we explored the oligomerization state of the enzyme in solution. Using size exclusion chromatography coupled to multi-angle light scattering (SEC-MALS), we showed that PPM1H^WT^-LD is a dimer (Fig 3B). In order to determine whether the contacts observed in crystals enable dimerization in solution, residues Gly357 and Ala359 were mutated to glutamate residues. The double mutant PPM1H^2Glu^-LD would be predicted to introduce longer side chains and negative charges that disrupt packing against α2 of the partner PPM1H molecule (Fig 3A). SEC-MALS analyses of PPM1H^2Glu^-LD revealed that it is indeed a monomer as designed (Fig 3B). The apparent physiological dimer can accommodate two Rab substrates without steric conflicts (Fig 3A). A potential PPM1H:pRab8A heterotetrameric complex would be organized with a two-fold symmetry axis parallel to the α3 helices in the flap domain (Fig 3A).

**Figure 3:**
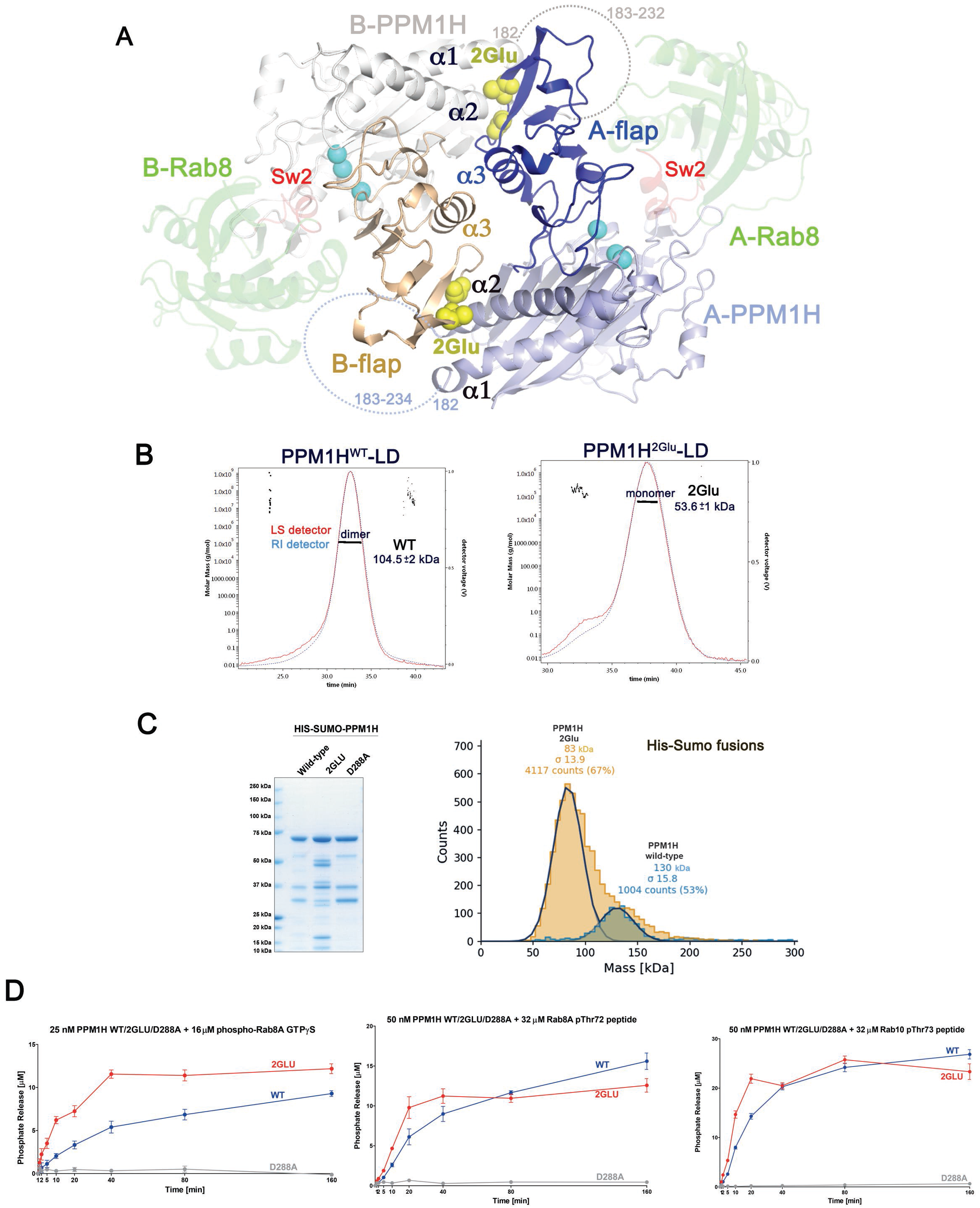
PPM1H is a dimer. (A) Model of the dimeric form of MnPPM1H^WT^-LD. Each monomer of PPM1H has an A or B prefix. Flap domains mediate the dimer and are shown in dark blue/wheat colors. The view is down a pseudo two-fold formed by α3 in the flap domain. Dotted lines denote the flexible loop (183-235) that connects α1/α2 in the catalytic domain. The dimeric organization of PPM1H positions this loop next to substrates, shown as docked models of pRab8a in green ribbons (transparent). The phosphoryated switch 2 helix of pRab8a is red. Yellow spheres indicate site of a double mutation in the flap domain (G357E+A359E) that disrupts a contact with α2 of the catalytic domain. **(B)** SEC-MALS analyses of PPM1H^WT^-LD and PPM1H^2Glu^-LD, showing that the WT enzyme is a dimer (104.5 kDa) while the double mutant is a monomer (53.6 kDa). The calculated molecular weight of His_6_-tagged PPM1H^WT^-LD used for the experiment is approximately 52.1 kDa. **(C)** His_6_-SUMO-tagged full-length variants of PPM1H were used for catalytic assays (*left panel*). 3 µg of recombinant wild-type or indicated mutant PPM1H proteins were resolved on 4-12% Bis-Tris gradient gel and stained with Instant Blue Coomassie. *Right*, mass photometry histogram for 40nM His_6_-SUMO-PPM1H WT (blue) and His_6_-SUMO-PPM1H 2Glu (brown), where WT is 130kDa (±15.8 kDa, with 1004 single molecules counted) and 2Glu is 83kDa (±13.9 kDa, with 4117 single molecules counted). The calculated molecular weight of the fusion protein is approximately 68.5 kDa. **(D)** *In vitro* malachite green assay time course of recombinant His_6_-SUMO-tagged PPM1H wild-type (blue), 2Glu (red), or D288A (grey) against 16 µM pThr72 phosphorylated Rab8a protein (GTPγS, *left*), 32 µM pThr72 phosphorylated Rab8a peptide (*middle),* or 32 µM pThr73 phosphorylated Rab10 peptide (*right*).

Full length PPM1H proteins fused to His_6_-SUMO at their N-termini were generated to probe whether the dimer is important for catalytic activity *in vitro*. The newly developed method of mass photometry was initially used to determine the oligomeric state of purified proteins in solution[31]. His_6_-SUMO-PPM1H^WT^ was observed to be a dimer at 40nM concentration, while the His_6_-SUMO-PPM1H^2Glu^ variant (in which residues Gly357 and Ala359 in the flap domain were both mutated to Glu) was a monomer (Fig 3C). The enzymatic activity of the monomeric variant of PPM1H was evaluated using pRab8a(GTPγS) and phospho-peptides from Rab8a and Rab10 (Fig 3D). Relative to WT, the monomeric variant of PPM1H appeared to be moderately more active against pRab8a protein. However, the catalytic activity of monomeric PPM1H was indistinguishable from the dimer using peptide substrates (Fig 3D). Therefore, the monomeric enzyme is active and dimerization is not required for efficient catalytic activity *in vitro*.

### PPM1J chimeras with the PPM1H flap domain are active against pRab8a

Although PPM1J is a close relative of PPM1H, previous work revealed negligible activity towards LRRK2 phosphorylated Rabs in cellular or biochemical assays[14]. Ignoring loops that are predicted to be unstructured and distant from the active site cleft, PPM1H and PPM1J have an identical domain organization (Figs 4A, EV4A). Intriguingly, residues in the catalytic domain of PPM1J that would face toward the substrate cleft are highly conserved and unlikely to account for specificity (Fig EV4B). The flap domains of PPM1H/J have relatively high sequence identities, but several sites that face the catalytic cleft are divergent (Figs EV4B, EV4C). To broadly explore the determinants of specificity, we engineered PPM1J chimeras that adopt the PPM1H anchor and/or flap domain (Figs 4A, 4B). The sequences of the chimeric proteins are provided in Zenodo repository file (dx.doi.org/10.5281/zenodo.5045023). The structure of PPM1H was critical for guiding the sites for grafts and thereby maintaining the integrity of the core catalytic fold. These recombinant proteins were successfully purified, and their catalytic properties were evaluated. Strikingly, the PPM1J chimeric proteins adopting the PPM1H flap domain were markedly active towards the pRab8a protein displaying similar or even moderately enhanced activity than PPM1H (Fig 4C). We also noted that the PPM1J chimera proteins containing the PPM1H flap domain restored activity towards the pRab8a peptide substrates (Fig 4D). This indicates that a portion of the PPM1H flap domain is also contributing to the interaction with the peptide. PPM1J displayed detectable but low activity towards the Thr72-phosphorylated Rab8a peptide. In contrast, the PPM1J chimera possessing the PPM1H N-terminal anchor gained a moderate level of activity against both pRab8a peptide and protein substrates, suggesting a minor contribution of the PPM1H anchor toward pRab8a specificity. Full activity for the J/H double chimera also suggests that the anchor regions of PPM1H/J can be swapped without effects on catalysis and that the flap domain is the dominant factor in specificity. Grafting of the PPM1J flap onto PPM1H (H/J flap) abolished the catalytic activity against pRab8a peptide and protein substrates (Figs 4C, 4D). We also analysed the activity of chimeras using Thr73-phosphorylated Rab10 peptides (Fig 4D, *right panel*). The peptide mimic of pRab10 is more promiscuous since it is dephosphorylated by PPM1J, albeit at a lower rate than PPM1H. Here again substitutions with the PPM1H flap/anchor motif stimulated the rates at which PPM1J dephosphorylated the pRab10 peptide. We also analysed the activity of PPM1M in these assays and observed that PPM1M displays low but detectable activity towards pRab8a GTPγS as well as the pRab8a and pRab10 phospho-peptide substrates (Fig 3D).

**Figure 4:**
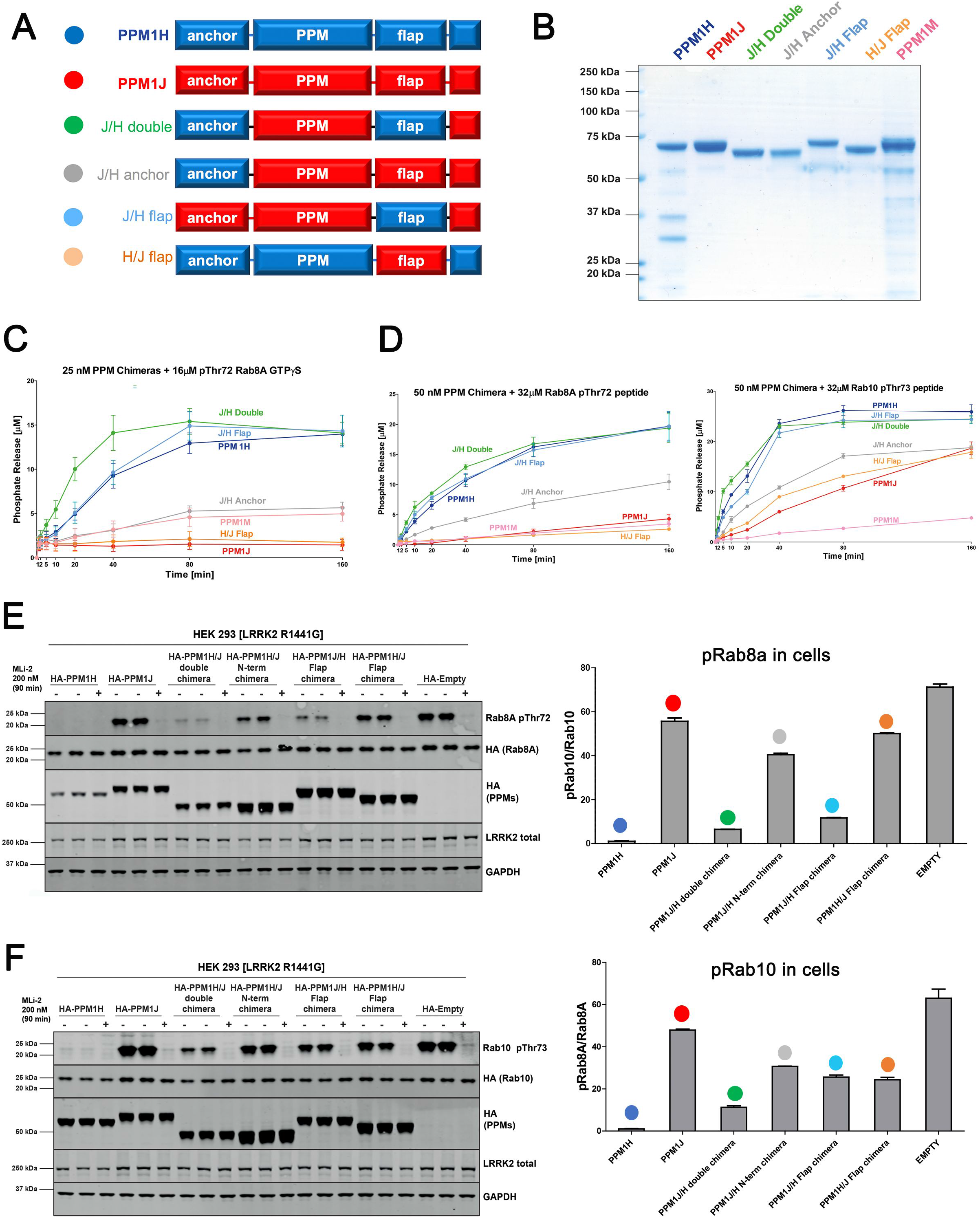
The flap domain of PPM1H is a determinant of Rab specificity. **(A)** Domain organization of chimeric variants for *in vitro* and cellular assays. **(B)** 3 µg of recombinant proteins were resolved on 4-12% Bis-Tris gradient gel and stained with Instant Blue Coomassie. **(C)** *In vitro* malachite green assay time course to determine enzyme activity of recombinant PPM proteins (25 nM for the protein substrate, 50 nM for the peptide substrate) against 16 µM pThr72 phosphorylated Rab8a protein (GTPγS). **(D)** *In vitro* malachite green assay against 32 µM pThr72-Rab8a peptide (*left*) and pThr73-Rab10 peptide mimic (*right*). **(E)** HEK293 cells overexpressing indicated constructs were treated with ± 200 nM MLi-2 for 90 min and then lysed. 10 µg whole cell lysate was subjected to immunoblot analysis with the indicated antibodies at 1 µg/ml final concentration and membranes were analysed using the OdysseyClx Western Blot imaging system. *Left*, each lane represents cell extract obtained from a different dish of cells (two replicates per condition without MLi-2 treatment, one replicate per condition with MLi-2 treatment). *Right*, the ratio of phospho-Rab8a/total Rab8a was quantified using Image Studio software and data presented relative to the phosphorylation ratio observed in PPM1H wild-type expressing cells. **(F)** As in **(E)** assessing phospho-Rab10 levels.

We further explored the activity of chimeric proteins in cellular assays. Lysates of HEK293 cells overexpressing pathogenic LRRK2[R1441G], Rab8a, and wild type or mutant forms of PPM1H were immunoblotted using pThr72-Rab8a phospho-specific antibody (Fig 4E). These assays reveal that PPM1J chimeras that adopt the PPM1H flap domain, in contrast to wild type PPM1J, dephosphorylate pRab8a to nearly the same extent as wild type PPM1H (Fig 4E). We also repeated these experiments with Rab10 instead of Rab8a and found that the PPM1J chimeras that adopt the PPM1H flap domain partially dephosphorylate Rab10, but to an intermediate level compared to PPM1H cells (Fig 4F). Overall, these results suggest that for cellular assays the gain of function by introducing PPM1H anchor and flap domains into PPM1J is less striking than *in vitro* assays. It is possible that additional factors such as cellular localization of PPM1H mutants regulate catalytic function in cells. Grafting the flap domain of PPM1J onto PPM1H resulted in marked loss of function in cellular assays of pRab8a and pRab10 hydrolysis (Figs 4E, 4F). Altogether, the *in vitro* and cellular assays suggest that the flap domain of PPM1H is an important determinant of substrate specificity for Rab GTPases.

### Mutagenesis and enzymatic assays of PPM1H

From the crosslinking and modelling studies, we identified Arg338 is a potential site of pRab8a binding despite being situated 27 Å from the catalytic ions (Fig 5A). In addition, a loop in the flap domain (386-396), with Leu392 at its center, forms an interface with α2 and the β-sandwich of the catalytic domain (Met252, Ala271, Cys269, Ile256). Despite the diversity of flap conformations among the PPM family, a short motif in this loop is highly conserved in sequence and structure, therefore it was targeted for mutagenesis. Finally, a lysine residue (Lys88) was identified, which is conserved as R33 in PPM1A (Fig EV1B) and is predicted to bind directly to the phosphate moiety of substrates[9]. Thus, three distinct sites were initial targets for mutagenesis and subsequent functional assays.

**Figure 5:**
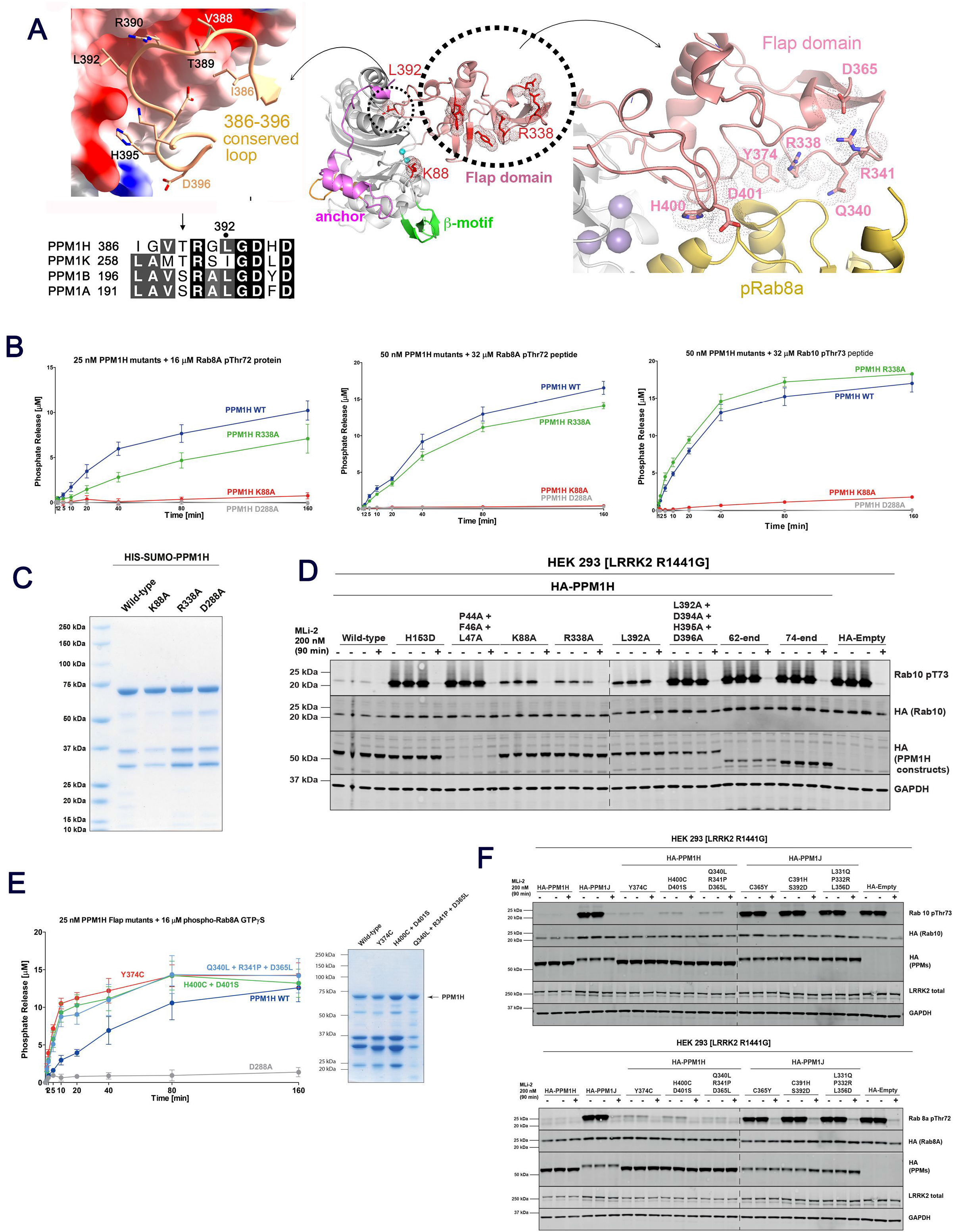
Mutagenesis and functional assays identify determinants of catalysis. **(A)** Sites for targeted mutagenesis on the flap domain and active site Lys88 residue. The central ribbon model provides context for the conserved loop (*left*) and epitopes distant from the active site in the flap domain (*right*). Residues subjected to mutagenesis are shown as stick models, with dotted envelopes in the central panel for emphasis. Docked pRab8a (yellow ribbons, *right*) highlights its proximity to mutagenized residues in the flap domain. **(B)** *In vitro* malachite green assay time course of recombinant PPM1H mutants against 16 µM pThr72 phosphorylated Rab8a protein (GTPγS, *left*), 32 µM pThr72 phosphorylated Rab8a peptide (*middle),* or 32 µM pThr73 phosphorylated Rab10 peptide (*right*). **(C)** 3 µg of recombinant wild-type or indicated mutant PPM1H proteins were resolved on 4-12% Bis-Tris gradient gel and stained with Instant Blue Coomassie (*left).* **(D)** HEK293 cells overexpressing indicated constructs were treated and analysed as described in Figure 4E. **(E)** Activity of 25 nM PPM1H flap domain mutants against 16 µM pThr72 phosphorylated Rab8a protein (GTPγS) using the malachite green assay time course. Quality of purified proteins (*right panel*) is shown using the protocol in **(C)**. **(F)** HEK293 cells overexpressing indicated constructs were treated and analysed as described in **(D)**.

We firstly evaluated the impact of the K88A and R338A mutations in a quantitative biochemical time course assay using either pThr72-Rab8a complexed to GTPγS as a substrate (Fig 5B, left) or pRab8a/pRab10 phospho-peptide substrates (Fig 5B, *middle/right*). For these assays, full-length PPM1H variants with an N-terminal SUMO tag were purified (Fig 5C). Although there were impurities in these preparations, their levels relative to intact enzyme were similar among the WT and mutants. The exception was L392A whose low yield of intact enzyme precluded quantitative analyses. The assays revealed that the PPM1H[K88A] mutation reduced initial rate activity by over 20-fold using both protein and phospho-peptide substrates. This is consistent with Lys88 playing a key role in mediating a direct contact with phospho-threonine of substrates. The R338A mutation reduced PPM1H initial activity approximately 2-fold when assayed with the pRab8a protein (Fig 5B, *left*) but had no significant impact when assayed with pRab8a/10 peptides (Fig 5B, *middle/right*). This result is consistent with Arg338 in the flap domain contributing towards optimal docking of pRab8a protein substrate, but not the peptide.

Using the HEK293 cell assay, we next investigated the role of the conserved loop from the flap domain, which forms an interface with the catalytic core (Fig 5D). The L392A mutant was expressed at 2 to 3-fold lower levels compared to wild type and displayed reduced activity towards phosphorylated Rab10. We also generated a quadruple mutant (L392A+D394A+H395A+D396A) that eliminates several interactions simultaneously between the loop and the catalytic domain. When expressed in HEK293 cells, this quadruple mutant lacked detectable activity towards phosphorylated Rab10 and was expressed ∼3-fold lower levels than wild type PPM1H (Fig 5D). Therefore, this conserved loop in the flap domain likely contributes to folding and may also affect catalysis. We also assessed the impact of K88A and R338A mutations in the HEK293 cell assay and observed that both mutations were expressed at comparable levels to wild type and displayed reduced ability to dephosphorylate pRab10. The significance of the mutant P44A+F46A+L47A (Fig 5D) is discussed below in the context of the ‘anchor’ region of PPM1H.

In addition to the above mutations, we identified potential Rab-interacting sites in the flap domain that are not conserved in PPM1J (Figs EV4C, 5A). We generated three mutants of PPM1H – Y374C, H400C+D401S, Q340L+R341P+D365L – which converted PPM1H residues to their PPM1J counterparts in the flap domain. Although the WT and recombinant proteins show partial degradation, they all retain catalytic activity against pRab8a (Fig 5E). A corresponding cellular assay involving these mutants similarly revealed no significant reduction in catalytic activity against either pRab8a or pRab10 substrates (Fig 5F). The reverse mutations in the flap domain (C365Y, C391H+S392D, L331Q+P332R+L356D) were also generated on a PPM1J background to see whether these sites enabled a gain-of-function phenotype. However, cellular assays revealed no significant activity against pRab8a and pRab10 (Fig 5F). These observations suggest that the specificity of PPM1H for Rab substrates does not localize to a single dominant epitope in the flap domain.

### Anchor-like folding motif in the PPM1H/J/M subfamily

PPM1H possesses a novel N-terminal extension preceding the core catalytic domain (Fig 6A). S33 interacts with the C-terminus of α2, adjacent to the flap domain, before winding around the back of the enzyme to the opposite side of the β-sandwich. PPM1A and PDP1 have an additional β-strand (β1) at the edge of the first β-sheet. In PPM1H, part of this anti-parallel β1 strand is conserved, while the remainder is dislodged and forms a loop preceding a shortened β2 strand. This distinct β1 strand conformation of PPM1H is due to the presence of a short 3_10_ helix (residues 43-47) in the N-terminal extension. The 3_10_ helix facilitates the insertion of the aromatic ring of F46 into the hydrophobic core of the β-sandwich (Fig 6A). Resembling an ‘anchor’, the side chains of P44, F46, and L47 cap the hydrophobic core of the β-barrel, effectively substituting for strand β1. To probe its significance, systematic deletion of the N-terminal residues of PPM1H toward the 3_10_ helix was performed (Fig 6B). Strikingly, PPM1H is relatively active until R43 is deleted (construct 44-end), which leads to reduced expression in HEK293 cells and loss of activity towards LRRK2 phosphorylated Rab10. To probe the importance of residues comprising the 3_10_ helix we mutated P44, F46 and L47 to alanine and observed that this nearly abolished PPM1H soluble expression in HEK293 cells, emphasizing their importance for enzyme folding (Fig 5D). Immediately following the 3_10_ helix, an α-helix (αN, N51-A58) and loop (D59-I65) interact directly with an active site loop (V83-D94).

**Figure 6:**
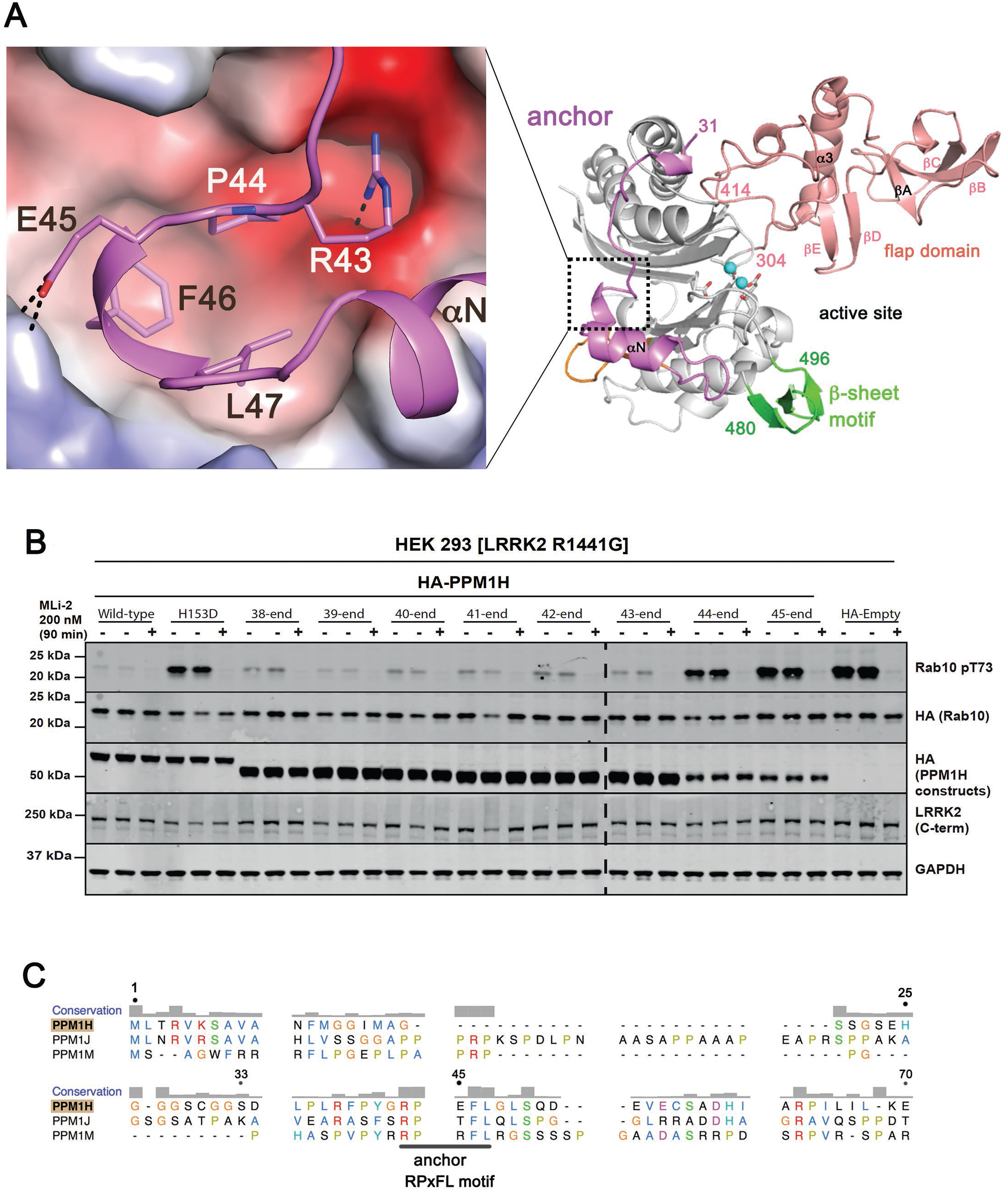
Anchor of PPM1H is a folding motif. **(A)** Interactions between the anchor of PPM1H against the electrostatic surface of the core catalytic domain. The region is placed in context with the dotted box on a ribbon model of PPM1H (*right*). **(B)** Incremental deletions of the N-terminus from residues 1-37 to 1-44, one residue at a time. Upon deletion of R43 (44-end), a reduced level of soluble PPM1H expression has no significant catalytic activity. **(C)** Sequence alignment of the N-terminal regions of the evolutionarily related PPM1H/J/M enzymes. The degree of conservation is above the alignment, and residue numbers correspond to PPM1H. A conserved anchor motif (RPxFL) is annotated below the sequences.

Despite sequence diversity at their N-termini, the PPM1H/J/M subfamily of phosphatases have a conserved RPxFL motif (Fig 6C). This anchor motif is likely involved in the folding of the catalytic domain through hydrophobic interactions with the β-sandwich. It may also influence catalysis due to its protrusion against the active site and a tether-like connection between the flap domain and the β-motif.

### Model for PPM1H specificity for Rab GTPases

The PPM family of phosphatases have a common catalytic domain with the incorporation of diverse structural elements that facilitate their specificity and function. The flap is adjacent to the active site and poorly conserved in sequence and structure within the PPM family. It has been associated with substrate specificity and catalysis[13, 32], but previous chimeras involving the flap region have been unsuccessful in transferring functions [10]. Here we provide evidence that PPM1H phosphatase specificity for Rab GTPases is encoded by the flap domain. Substitution of this domain into PPM1J leads to its ability to hydrolyze phospho-Rab8a both *in vitro* and in cellular assays. Also, we have identified a highly conserved loop motif from the flap domain (residues 386-396) that appears to contribute to folding and/or catalysis. This conserved loop in human PPMs, with Leu392 at its center (PPM1H numbering), has not been investigated previously. Although speculative, we think it might couple substrate binding to catalysis given its proximity to the active site. Intriguingly, the equivalent Leu392 hydrophobic pocket has recently been identified as a ‘regulatory switch’ in bacterial phosphatases[33]. Finally, an anchor preceding the catalytic domain caps the hydrophobic core of the β-sandwich and has apparently evolved as a folding motif in the PPM1H/J/M subfamily. In a PPM1J chimera, the PPM1H anchor and flap domain act in concert to promote the dephosphorylation of pRab8a and pRab10. A cartoon model depicting the structural attributes that encode PPM1H specificity for phospho-Rab GTPases is shown in Fig 7. Although extended flap domains are conserved in evolutionarily related PPM1J and PPM1M, differences in their sequences and/or conformations likely encode the specificity of PPM1H for phosphorylated Rab8a/10. Further details of PPM1H specificity await the structure of a substrate-trapped complex.

**Figure 7:**
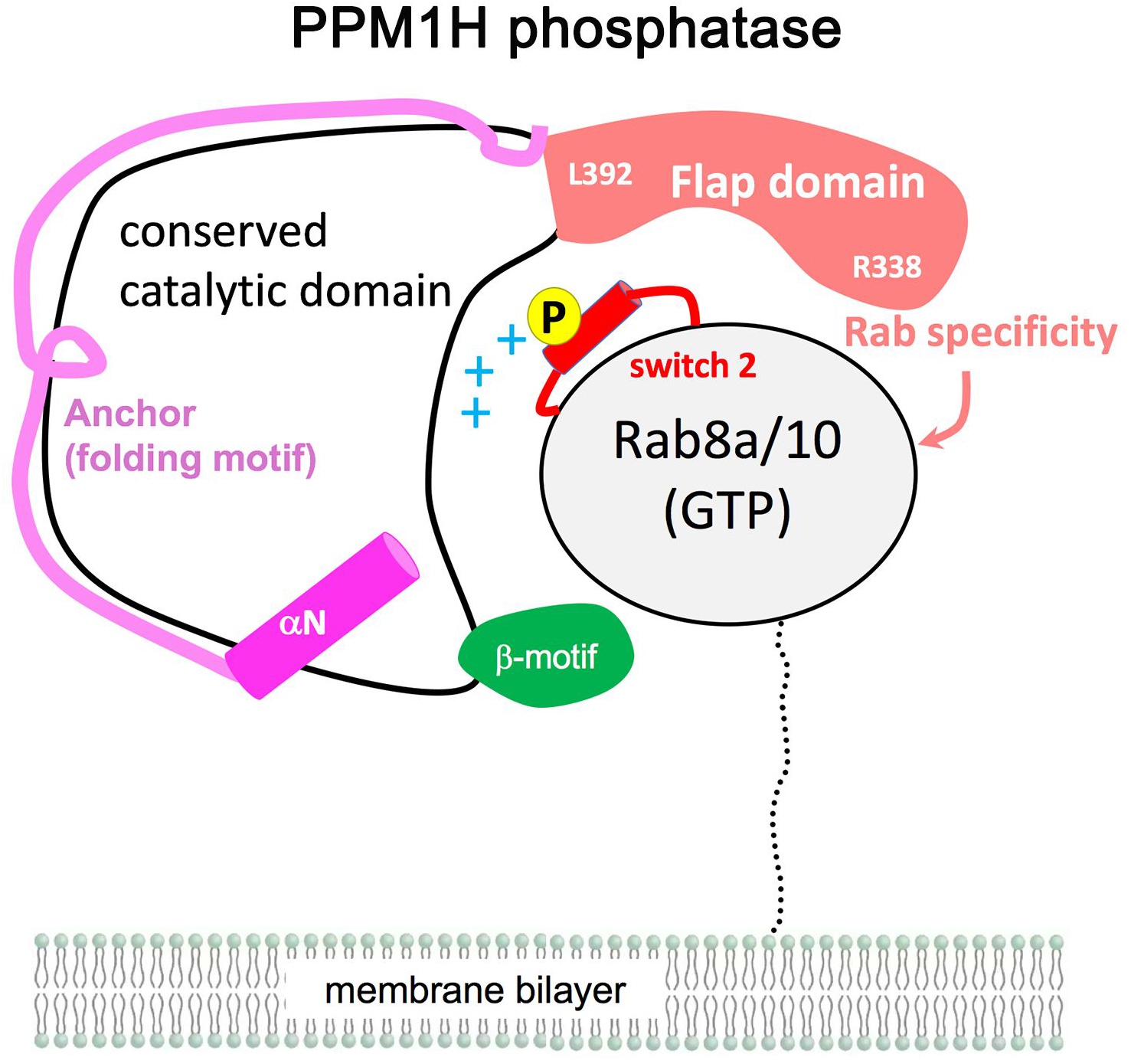
Model of PPM1H specificity for Rab GTPases. Although PPM1H phosphatase is likely to be dimeric in cells, a 1:1 complex is shown for simplicity. The switch 2 helix is red and the phosphate site is yellow. The flap domain (305-414) encodes the primary specificity for phosphorylated Rabs while the anchor (33-79) contributes to folding of the enzyme. The β-motif (green, residues 480-496) is also indicated. The conserved loop motif (386-396) with a central Leu392 residue may couple substrate binding to catalysis.

We also find that PPM1H is a physiological dimer in solution *via* contacts mediated by the flap domain. DSBU-induced crosslinks between a flexible loop 183-235 and pRab8a can be attributed to dimerization. This loop is distant from the active site, but in a docked model of a heterotetrameric enzyme:substrate complex, the loop is situated in close proximity to pRab8a (Figs 2B, 3A). At least one of the residues in this loop (Ser210) is phosphorylated [34] and in future work it would be important to investigate the role that dimerization plays in controlling PPM1H function in cells. However, conversion of the enzyme into a monomer by mutagenesis through abolition of flap/α2 contacts does not markedly affect catalysis *in vitro*. PPM1H has been localized to Golgi, where it likely counters the LRRK2 pathway by dephosphorylating Rab8a and Rab10 under physiological conditions[14]. Rab proteins have a flexible stretch of ≈30 residues between their G domain and their prenylated C-terminal tails that localize them to subcellular membranes (Fig 7). In future work it would be important to dissect the mechanism by which PPM1H localizes to membranes to hydrolyze phophorylated Rab substrates.

## Materials and Methods

### Constructs and expression of proteins

The cDNA for PPM1H^DA^-LD (residues 33-514, D288A) with the residues 188-226 replaced by a short GSGS motif was synthesized as a codon-optimized gene for *E. coli* expression (Genscript, Inc). The cDNA was cloned into pET28a-(+)-TEV at the NdeI/BamH1 restriction sites. The PPM1H^WT^-LD construct was made by site directed mutagenesis using the following primers: 5’-GTG GCG AAC GCG GGT GAT AGC CGT GCG ATC ATT ATC-3’ (for) and: 5’-GAT AAT GAT CGC ACG GCT ATC ACC CGC GTT CGC CAC-3’ (rev). Full-length and 57-end cDNA variants of PPM1J were obtained from Genscript with codon-optimization for *E.coli* expression. Using NdeI and BamH1 sites at their 5’ and 3’ ends, the genes were inserted into the same vector above. The PPM1H^DA^ (residues 33-514) construct was amplified using the following primers: 5’-TACTTCCAATCC <u>TCG</u> GAC CTG CCC CTG CGT TTC3’ (for) and 5’-TATCCACCTTTACTG <u>TTA TCA</u> TGA CAG CTT GTT TCC ATG-3’ (rev); S33(PPM1H)/stop codons underlined. We used a human PPM1H (NM_020700) template carrying the D288A mutation and LIC cloned the resulting DNA into the pNIC28-BSA4 vector which allows the expression of a hexahistidine -tagged protein. All plasmids generated by PCR were confirmed by sequencing. For the full-length PPM1H PhosTag assay, residues 1-514 of PPM1H were LIC cloned into the pLIC-MBP vector[35]. The primers used to amplify PPM1H from the template pET15b-Sumo-PPM1H (University Dundee DU62790) were: 5’-C CAG GGA GCA GCC TCG ATG CTC ACT CGA GTG AAA TC-3’ (for) and 5’-GC AAA GCA CCG GCC TCG TTA TCA TGA CAG CTT GTT TCC-3’ (rev).

Expression of PPM1H was carried out in LB Broth supplemented with 30 µg/ml kanamycin (FORMEDIUM™). After incubation at 37°C to an OD_600_ of ∼0.6 the culture was cooled down to 18°C and induced with 0.5 mM IPTG (FORMEDIUM™), after which cells were grown at 18°C overnight. Cells were harvested by centrifugation and the pellets were resuspended in His-tag extraction buffer (20 mM Tris-HCl pH 8.0, 300 mM NaCl, 5 mM MgCl_2_, 20 mM imidazole and 10 mM β-mercaptoethanol). Purification of PPM1H in complex with Mn^2+^ involved substitution of MgCl_2_ with MnCl_2_ in all steps. After lysis by sonication the cell lysate was centrifuged at 26,000 x g for 45 minutes at 4°C to remove cellular debris. The supernatants were loaded onto a nickel agarose resin (QIAGEN) in a gravity flow setup. The resin was washed with a 10-fold excess of extraction buffer and 5-fold excess wash buffer (extraction buffer supplemented with 40 mM Imidazole). The hexahistidine-tagged protein was then eluted using extraction buffer supplemented with 200 mM imidazole.

In the case of proteins for crystallization removal of the His_6_ tag was performed by overnight incubation at 4°C in dialysis against gel filtration buffer (20mM Tris-HCl pH 8.0, 100mM NaCl, 5mM MgCl_2_, 1mM DTT) using recombinant TEV protease, followed by a second Ni^2+^-agarose column. The ‘flow-through’ fractions were collected, while the uncut proteins remained on the resin. Soluble aggregates were eliminated by running the sample through a Superdex 200 (10/300) gel filtration column (GE Healthcare) equilibrated in gel filtration buffer. The peak fractions of PPM1H were pooled and concentrated in 10kDa MWCO concentrator tubes prior to crystallization trials. A detailed protocol describing the expression and purification of PPM1H has been reported (dx.doi.org/10.17504/protocols.io.bu7wnzpe).

### Phosphorylation of Rab8a Q67L and Rab10 Q68L

The *in vitro* phosphorylation of Rab8a (T72) by MST3 kinase and subsequent purification has been described previously[28] and a detailed protocols.io protocol reported (<u>dx.doi.org/10.17504/protocols.io.butinwke</u>). In brief, Rab8a was mixed with MST3 at a 8:1 molar ratio, and incubated overnight. The phosphorylation buffer was adjusted to the following conditions: 50mM Tris-HCl, 150mM NaCl, 10mM MgCl_2_, 2mM ATP, pH 7.5. The phosphorylation mixture was then dialyzed against low salt buffer and loaded to a MonoS (GE HEalthcare) column. Phosphorylated Rab8a was separated from unphosphorylated by ion exchange chromatography applying a 50% gradient low to high salt buffer (10mM MES, 10mM (low) or 1M NaCl (high), 5mM MgCl_2_, 1mM DTT, pH 5.2). The phosphorylation of Rab8a was confirmed by PhosTag gel electrophoresis prior to subsequent experiments. pRab10 was prepared by phosphorylating Rab10 (1-181) by MST3 using a similar procedure described previously (<u>dx.doi.org/10.17504/protocols.io.bvjxn4pn</u>).

### Crystallization, data collection and refinement

Crystals of PPM1H variants were grown at concentrations between 5-10 mg/L using the vapour diffusion method. Crystals were harvested in precipitant supplemented with 30% glycerol and stored frozen in liquid nitrogen. X-ray data were collected under a cryogenic nitrogen stream (100K) at the FMX beamline, NSLSII synchrotron (Brookhaven, New York, USA). Details of crystallization conditions and the quality of diffraction data are shown in Table 1. Native diffraction data were reduced using XDS and Aimless, followed by structure determination using the Phaser software in the PHENIX package[35, 36]. Molecular replacement using Phaser (PPM1A; PDB code 6b67[9]) was successful using data from crystals of PPM1H^DA^-LD, and subsequently, Autobuild in Phenix provided an initial model for PPM1H. Refinement was performed through multiple rounds of energy minimization and model building using Phenix and Coot software[37]. The structures of the other variants of PPM1H (PPM1H^WT^, PPM1H^WT^-LD, MnPPM1H^WT^-LD) were determined using PPM1H^DA^-LD as a search model and subsequently refined using Phenix and Coot. These higher resolution models were useful for the final refinement of PPM1H^WT^-LD. To reduce model bias, all four structures have common reflections flagged for the R-free data set. PPM1H^WT^-LD, PPM1H^DA^-LD and MnPPM1H^WT^-LD have a single cysteine mutation (C56A) at a non-conserved residue that enhances crystallizability. The asymmetric unit for all structures contains two molecules of PPM1H. Two non-native residues (His-Met) from the cleaved affinity tag are seen in one of the molecules of PPM1H^DA^-LD at its N-terminus. Statistics from data collection and refinement are shown in Table 1.

### Docking of PPM1H and pRab8a

Docking was performed using Haddock software [30] with distance restraints between the active site and pThr72 of pRab8a (PDB code 7lwb; [38]). Initial attempts at docking of PPM1H variants with two Mg^2+^ ions resulted in solutions with >7Å distance between the metal ions and the switch 2 phosphate of pRab8a. More realistic distances between the active site and pRab8a were achieved using the structure of MnPPM1H^WT^-LD, including a direct contact (3 Å) between the third metal site (M3) and pThr72. No other distance restraints between the enzyme and substrate were applied to enable an unbiased docking calculation. The structures were stripped of waters and refined with explicit solvent followed by default CNS scripts (Crystallography and NMR System) that performed semi-flexible simulated annealing and docking calculations. Docking of the switch 2 phosphopeptide (residues 65-79) was performed by extracting the coordinates from the structure of pRab8a. The N-terminus was capped by an acetyl group while the C-terminus was capped by an amide to eliminate charges. Docking calculations were performed using the same strategy outlined for the the G domain of pRab8a.

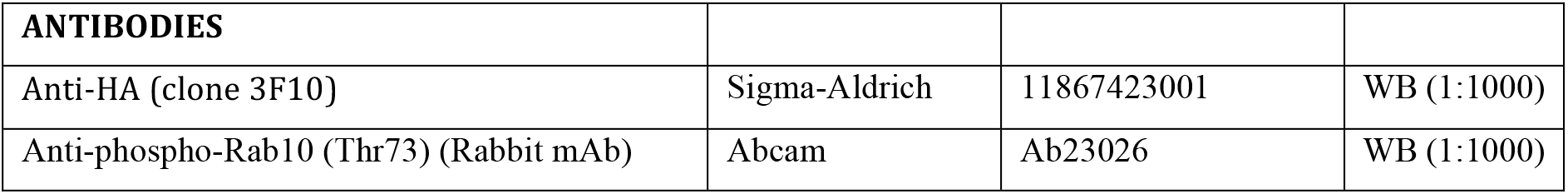
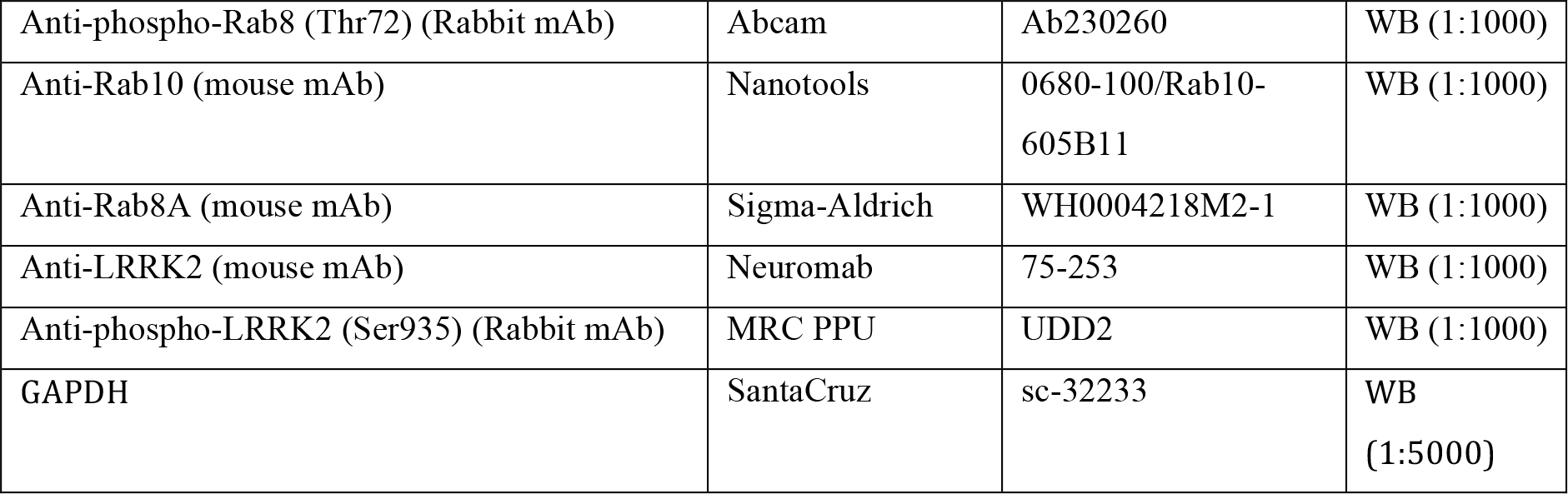

| **ANTIBODIES** |  |  |  |
| --- | --- | --- | --- |
| Anti-HA (clone 3F10) | Sigma-Aldrich | 11867423001 | WB (1:1000) |
| Anti-phospho-Rab10 (Thr73) (Rabbit mAb) | Abcam | Ab23026 | WB (1:1000) |
| Anti-phospho-Rab8 (Thr72) (Rabbit mAb) | Abcam | Ab230260 | WB (1:1000) |
| Anti-Rab10 (mouse mAb) | Nanotools | 0680-100/Rab10-605B11 | WB (1:1000) |
| Anti-Rab8A (mouse mAb) | Sigma-Aldrich | WH0004218M2-1 | WB (1:1000) |
| Anti-LRRK2 (mouse mAb) | Neuromab | 75-253 | WB (1:1000) |
| Anti-phospho-LRRK2 (Ser935) (Rabbit mAb) | MRC PPU | UDD2 | WB (1:1000) |
| GAPDH | SantaCruz | sc-32233 | WB<br>(1:5000) |

### Plasmids for expression in HEK293 cells

Expression constructs were obtained from the MRC-PPU repository at the University of Dundee: Detailed sequences of each construct is reported on the https://mrcppureagents.dundee.ac.uk/ website

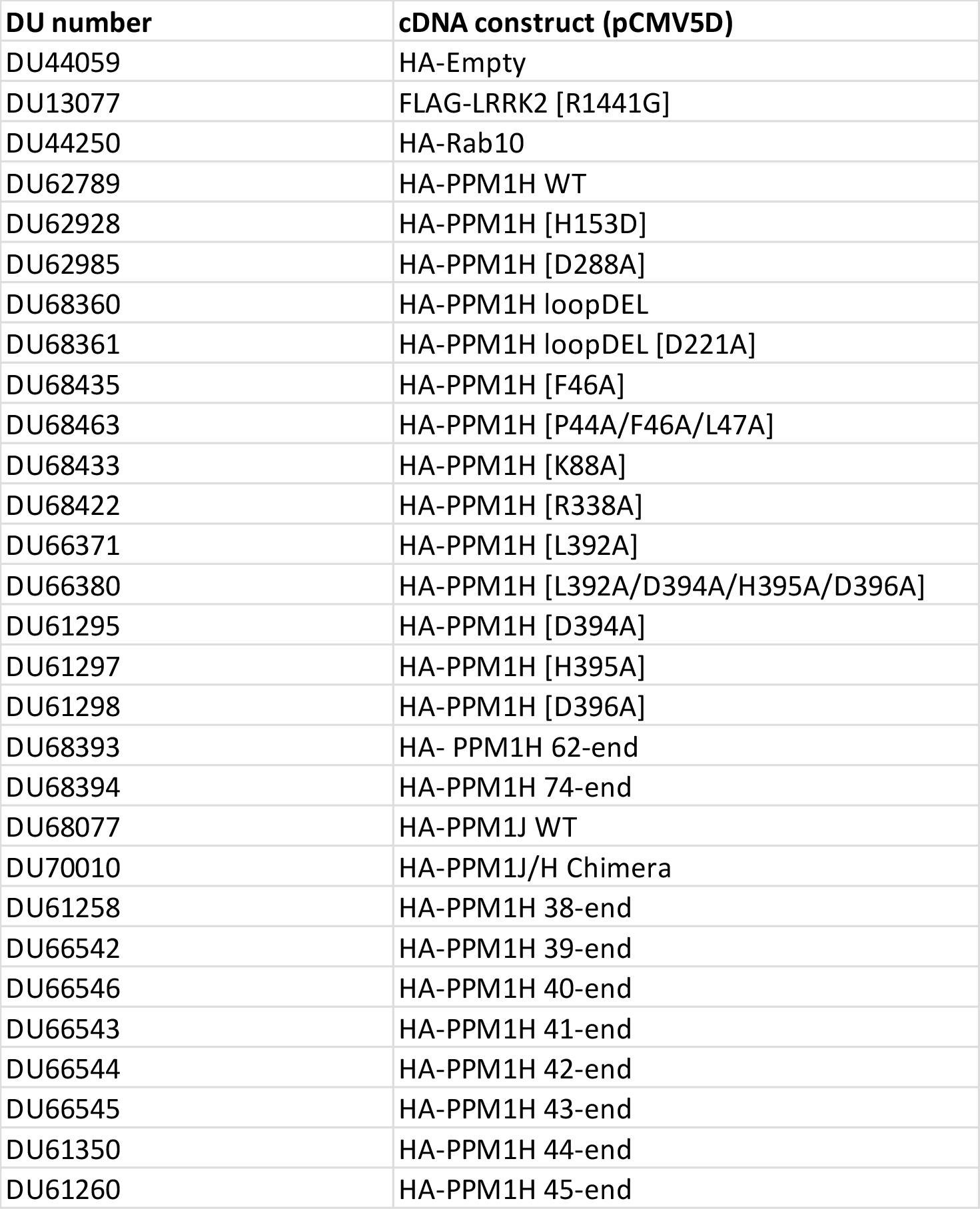
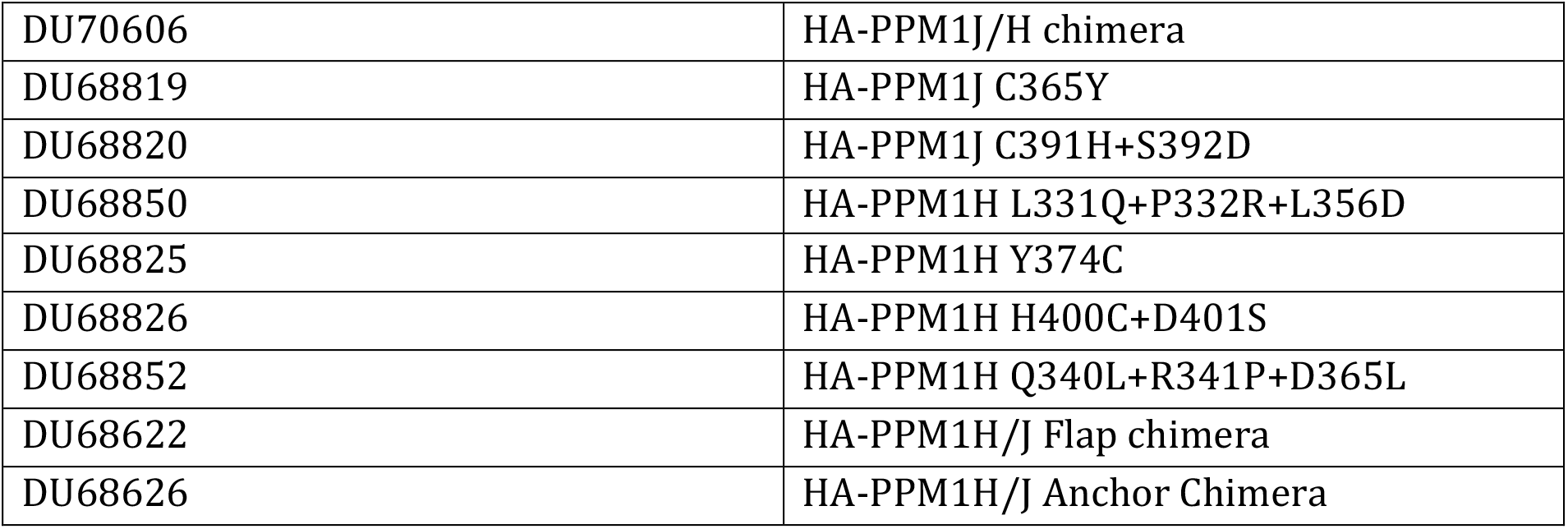

DNA constructs were amplified in *Escherichia coli* DH5α and purified using a Hi-Speed Plasmid Maxi Kit (Qiagen).

### Cell culture, transfections, treatment, and lysis

HEK293 cells were cultured in Dulbecco’s modified Eagle medium (Gibco) supplemented with 10% (v/v) foetal bovine serum, 100 U/ml penicillin and 100 µg/ml streptomycin at 37°C and 5% CO2. Cells used in this study were regularly tested for mycoplasma contamination. Transient transfections were performed 24 h prior to cell lysis using polyethylenimine PEIMax (0.1% w/v) (Polysciences)[39]. Cells were grown to ∼70% confluency in 6-well (3.5 cm well diameter) plates prior to transfection. 1.75 µg total plasmid DNA was mixed with 5.5 µl PEI in 150 µl Opti-MEM I Reduced Serum Media (Gibco) and incubated for 30 min at room temperature. The DNA-PEI mixture was added drop-wise to cells. Cells were treated ± MLi-2 LRRK2 inhibitors[40] at a final concentration of 200 nM 90 min prior to lysis. Cells were lysed in 150 µl ice-cold lysis buffer (50 mM Tris-HCl pH 7.5, 150 mM NaCl, 10% glycerol, 10 mM 2-glycerophosphate, 10mM sodium pyrophosphate, 1 mM sodium orthovanadate, 1 µg/ml microcystin-LR, complete EDTA-free protease inhibitor cocktail (Roche), and 1% (v/v) Triton X-100) and collected in 1.5 ml Eppendorf tubes. Lysates were clarified by centrifugation at 20 800 *g* at 4°C for 20 min and supernatants were quantified by Bradford assay before subjected to immunoblot analysis.

### Immunoblot analysis

Clarified cell or tissue extracts were mixed with a quarter of a volume of 4× SDS–PAGE loading buffer [250 mM Tris–HCl, pH 6.8, 8% (w/v) SDS, 40% (v/v) glycerol, 0.02% (w/v) Bromophenol Blue and 4% (v/v) 2-mercaptoethanol]. 10-30 µg of samples were loaded onto NuPAGE 4–12% Bis–Tris Midi Gel (Thermo Fisher Scientific, Cat# WG1403BOX) and electrophoresed at 130 V for 2 h with the NuPAGE MOPS SDS running buffer (Thermo Fisher Scientific, Cat# NP0001-02). At the end of electrophoresis, proteins were electrophoretically transferred onto the nitrocellulose membrane (GE Healthcare, Amersham Protran Supported 0.45 µm NC) at 100 V for 90 min on ice in the transfer buffer (48 mM Tris–HCl and 39 mM glycine). Transferred membrane was blocked with 5% (w/v) skim milk powder dissolved in TBS-T [20 mM Tris–HCl, pH 7.5, 150 mM NaCl and 0.1% (v/v) Tween 20] at room temperature for 1 h. The membrane was typically cropped into three pieces, namely the ‘top piece’ (from the top of the membrane to 100 kDa), the ‘middle piece’ (between 100 and 37 kDa) and the ‘bottom piece’ (from 37 kDa to the bottom of the membrane). The top piece was incubated with rabbit anti-LRRK2 pS935 UDD2 antibody multiplexed with mouse anti-LRRK2 C-terminus total antibody diluted in 5% (w/v) bovine serum albumin in TBS-T to a final concentration of 1 µg/ml for each of the antibody. The middle piece was incubated with rat anti-HA antibody diluted in 5% (w/v) bovine serum albumin in TBS-T to a final concentration of 100 ng/ml. The bottom pieces were incubated with mouse anti-GAPDH multiplexed with rabbit MJFF-pRab10-clone-1 monoclonal antibody and rat anti-HA antibody diluted in 5% (w/v) bovine serum albumin in TBS-T to a final concentration of 0.5 µg/ml for each of the antibody. All blots were incubated in primary antibody overnight at 4°C. Prior to secondary antibody incubation, membranes were washed three times with TBS-T for 10 min each. The top and bottom pieces were incubated with goat anti-mouse IRDye 680LT (#926-68020) secondary antibody multiplexed with goat anti-rabbit IRDye 800CW ((#926-32211) secondary antibody diluted in TBS-T (1:25, 000 dilution) for 1 h at room temperature. The middle piece was incubated with goat anti-mouse IRDye 800CW (#926-32210) secondary antibody diluted in TBS-T (1 : 25, 000 dilution) at room temperature for 1 h. Membranes were washed with TBS-T for three times with a 10 min incubation for each wash. Protein bands were acquired via near infrared fluorescent detection using the Odyssey CLx imaging system and quantified using the Image Studio software. A detailed protocols.io protocol for immunoblotting LRRK2 and pRabs has previously been described (dx.doi.org/10.17504/protocols.io.bsgrnbv6)

### *E.coli* expression and purification of recombinant phosphatases

Plasmids (DU62835; DU68087; DU68140; DU68554; DU68559; DU68560; DU68621; DU68625; *DU68875,* DU70009, DU68875, DU70609, DU68822, DU68823, DU68851, DU68828, DU68829, DU68853, DU68141 all available from MRC-PPU Reagents: mrcppureagents.dundee.ac.uk) were transformed into *Escherichia coli* BL21-DE3-pLysS and expressed in 3 x 1 litre of Lucia Broth medium (Merck) supplemented with 100µg/ml ampicillin (Formedium). Bacteria were cultured at 37°C until OD600 reached 0.8. The temperature was reduced to 16°C and after 1 h protein phosphatase expression was induced by addition of 25 µM Isopropyl β-D-1-thiogalactopyranoside for incubation overnight. Cells were harvested by centrifugation at 4200 x g for 30 min at 4°C before being resuspended in 20ml collection buffer (50 mM Tris-HCL pH 7.5, 150 mM NaCl, 20 mM imidazole, 2 mM MgCl_2_, 7 mM 2-mercaptoethanol). The resuspension was made to 5% (by vol) glycerol, 300 mM NaCl, 0.4% Triton X-100, 1 mM AEBFS (Pefabloc®), and 10 µg/ml Leupeptin. Cell suspension was sonicated, and insoluble material was removed by centrifugation at 40,000 x g at 4°C for 25 min. 2ml Cobalt resin was equilibrated in collection buffer before incubated with lysates for 2 h at 4°C. After incubation, resin was washed 5 times with 7 volumes wash buffer (50 mM Tris-HCL pH 7.5, 250 mM NaCl, 20 mM imidazole, 2 mM MgCl_2_, 7 mM 2-mercaptoethanol, 5% (v/v) glycerol). The resin was transferred onto a 5 ml polyprep (Biorad) filtration device. Protein was eluted with elution buffer (wash buffer diluted with 1 M imidazole to 0.4 M imidazole) and 1 ml fractions were collected. Protein-containing fractions were pooled and 1.6 ml protein was subjected to size exclusion chromatography using a Superdex 200 XK16/60 column (GE Healthcare) and equilibrated into 50 mM Tris-HCL pH 7.5, 200 mM NaCl, 2 mM MgCl_2_, 7 mM 2-mercaptoethanol, 0.015% (w/v) Brij35. Fractions containing PPM1H were pooled and concentrated before aliquoted and snap frozen to store at −80°C. A detailed protocols.io protocol for expression and purification of PPM1H has been reported (dx.doi.org/10.17504/protocols.io.bu7wnzpe)

### Quality control gel for recombinant phosphatases

3 µg of each protein was prepared in SDS-PAGE sample buffer and 5% 2-mercaptoethanol. Electrophoresis was carried out using NUPAGE Bis-Tris 4-12% gradient gels (Life Technologies) and run at 120 V. The gel was then stained for 1h using Instant Blue Coomassie (Expedeon). Protein concentrations were adjusted using the upper protein band, which corresponded to the undegraded protein.

### Quantitative phosphatase assays

The quantitatively Thr72 phosphorylated Rab8a complexed to either GTP or GDP was prepared as described previously[14]. The phospho-peptide encompassing the Thr72 LRRK2 phosphorylation site on human Rab8a (AGQERFRT*ITTAYYR residues 65 – 79, where T* - pThr) was synthesized and purified by JPT Peptide Technologies GmbH. Phosphate release from either the phospho-Rab8a protein or peptide was detected by using the malachite green phosphatase assay[41] according to the sigma protocol. The phosphatase assays were performed in 96-well flat-bottomed plates using a final volume of 80 µl and either 25 nM PPM phosphatase and 16 μM phospho-Rab8a protein or 50 nM PPM phosphatase and 32 μM phospho-Rab peptide. The phosphatase was diluted in 50 mM Tris-HCl pH 7.5, 200 mM NaCl, 2 mM MgCl_2_, 7 mM 2-mercaptoethanol, 5% (v/v) glycerol. The phospho-Rab8a protein and phospho-peptide are diluted in 40 mM HEPES buffer pH7.5, supplemented with 10 mM MgCl_2_ at a final reaction volume of 80 µl. The reactions were undertaken at room temperature and initiated by adding the diluted phosphatase to the phospho-Rab8 protein or phospho-peptide substrate. At the indicated times, the reactions were quenched by addition of 20 µl Malachite Green working reagent (Sigma Aldrich; MAK307). After incubation at room temperature for 30 min absorbance was measured at 620 nm. Free phosphate at increasing concentrations (1-40 µM) was used to make a standard curve, which was used to determine the concentration of phosphate released from each reaction. Malachite green working reagent and free phosphate standards were prepared fresh prior to each experiment, according to Sigma manual. Graphs were prepared using GraphPad Prism 5 software. To determine Km and Vmax of PPM1H against pRab8a protein and pRab8a/pRab10 peptide substrates, phosphatase reactions were performed as described above, except that the concentrations of phospho-Rab8a protein (complexed to either GDP or GTPγS) were varied between 2-32 μM, and for the pRab8a or pRab10 phospho-peptides between 2-256 μM. The initial rate (V_0_) was calculated by dividing concentration of released phosphate at 5 min by time and plotted against each substrate concentration. Enzyme kinetic constants were then obtained using GraphPad Prism 5 software. A detailed protocols.io protocol for quantifying PPM1H phosphatase activity towards LRRK2 phosphorylated Rab proteins and peptides using the Malachite Green method has previously been reported (dx.doi.org/10.17504/protocols.io.bustnwen).

### Phostag PPM1H/PPM1J phosphatase assay with phospho-Rab8a and phospho-Rab10

In vitro dephosphorylation assay was performed in a total volume of 20 µl in 40 mM Hepes buffer (pH 7.0) containing 10 mM MgCl_2_ using 2.5 µg pThr72 phosphorylated Rab8a or pThr73 phosphorylated Rab10 and increasing concentrations of the phosphatase. The assay was initiated by addition of PPM1H or PPM1J (3, 10, 30, 100, 300 ng) diluted into 40 mM HEPES Buffer pH 7.5 supplemented with 10 mM MgCl_2_. The assay was carried out for 20 min and terminated by addition of 4 x LDS (106 mM Tris HCl, 141 mM Tris Base, 2% (by mass) LDS, 10% (by vol) glycerol, 0.51 mM EDTA, 0.22 mM SERVA Blue G250, 0.175 mM Phenol Red, pH 8.5) with 5% (by vol) 2-mercaptoethanol. Samples were then subjected to Phos-tag gel electrophoresis to determine stoichiometry of phosphorylated Rab8a or Rab10 as described previously in [22]. Gel was stained using Instant Blue Coomassie (Expedeon).

### Mutagenesis, biophysical and functional studies of monomeric PPM1H

The cDNA for PPM1H^2Glu^-LD (residues 33-514, G357E+A359E) was synthesized as a codon-optimized gene for *E. coli* expression (Genscript, Inc). The protein sequence is identical to PPM1H^WT^-LD except for the double mutation that carries two glutamate residues in the flap domain. WT and mutant variants were purified as described above in extraction buffer supplemented with 5mM MgCl_2_. Following elution from Ni-agarose, the His_6_-tagged proteins were immediately purified on a Superdex 200 (10/300) column. The main peaks were collected for subequent characterization using a SEC-MALS system comprising an Agilent HPLC system coupled to a DAWN Heleos multi-angle light scattering system and an Optilab TrEX refractometer (Wyatt Corp). WT and mutant samples were injected at 2 mg/mL concentration (100μL) into a Superdex 200 column preceding the two detectors. Peaks were analyzed using Astra software (Wyatt Corp) to calculate the mass of PPM1H in solution. Astra data files have been uploaded to the Zenodo server (dx.doi.org/10.5281/zenodo.5045023).

### Crosslinking of PPM1H and pRab8a

Recombinant PPM1H[D288A] phosphatase (Purified as described in dx.doi.org/10.17504/protocols.io.bu7wnzpe) and recombinant stoichiometrically Thr72 phosphorylated Rab8a (purified as described in dx.doi.org/10.17504/protocols.io.butinwke) were used to perform the crosslinking analysis using DSBU crosslinker[24]. As the crosslinking reaction can be inhibited by amine containing buffers, a buffer exchange step was performed for both PPM1H and pRab8a protein solutions prior to the crosslinking. Proteins were be dialysed into 40 mM HEPES pH 7.5, 150 mM NaCl, 2 mM MgCl_2_ buffer using Diacon dialysis tubes (MD6-71, Molecular Dimensions). To form the complexes, a mixture of stoichiometrically Thr72 phosphorylated Rab8a at a concentration 35 µM (∼0.7 mg/ml, migrates at 20 kDa) and PPM1H at 28 µM (∼1.4 mg/ml, migrates at 50 kDa) was prepared in total volume of 15.5 µl using 40 mM HEPES pH 7.5, 150 mM NaCl, 2 mM MgCl_2_ as a dilution buffer (1.25-fold molar excess of Rab8a to PPM1H) and incubated at 30°C for 1 h, and cooled down to RT for 10 min. A fresh 300 mM stock solution of the crosslinker was prepared by dissolving 1 mg of DSBU in 15.5 µl of anhydrous DMSO. An aliquot of the crosslinker solution (0.5 µl) was added into 15.5 µl of the protein solution and mixed, which resulted in a final concentration of 9.375 mM of DSBU (Disuccinimidyl Dibutyric Urea, Thermo Scientific™, A35459) in 16 µl of reaction mix, giving a 250-fold molar excess to pRab8a and 335-fold molar excess to PPM1H. The crosslinking reaction was performed at RT for 10 min and quenched by addition of 2 µl of 1 M Tris pH 8.8, followed by addition of 7 µl of 4 x NuPAGE LDS sample buffer (without reducing agents). The samples were immediately resolved by SDS-PAGE using a 4-12% gradient NuPAGE Bis-Tris gel. The final reaction volume was 25 µl, which allowed to run 6 µl of the sample 4 times, to use 4 different digestion conditions. After the SDS-PAGE run, the gel was fixed and stained for 2 h using Invitrogen™ Colloidal Blue Staining Kit and de-stained in milli-Q water (following the kit manual).

Selected bands were excised from the gel, cut into 1 mm cubes, and placed in low-binding 1.5 ml Eppendorf tubes. Gel pieces were dehydrated by addition of 500 µl of acetonitrile (ACN) and incubation for 10 minutes at RT. Supernatant was discarded and the samples were reduced by adding 50 µl of freshly prepared 10 mM DTT solution and incubating at 56°C for 30 min with mixing (1200 rpm). After this incubation 500 µl of ACN was added and the samples incubated at RT for 10 minutes. Supernatant was discarded and the samples were alkylated by adding 50 µl of freshly prepared 55 mM iodoacetamide (IAA) solution and incubating at RT for 20 min with mixing (1200 rpm). After this incubation 500 µl of acetonitrile was added and the samples incubated at RT for 10 minutes. Supernatant was discarded and 100 µl of 50 mM ABC in 50% (v/v) acetonitrile was added and the samples incubated at RT for 30 min. After this incubation 500 µl of acetonitrile was added and the samples incubated at RT for 10 minutes, supernatant was discarded. Fresh solutions of proteases were prepared: trypsin at a concentration of 50 ng/µl, AspN at a concentration of 20 ng/µl, GluC at a concentration of 50 ng/µl. 20 µl of Trypsin solution was added to the dehydrated gel pieces. For the double-digestion conditions 20 µl of AspN solution or 20 µl of GluC solution was added following the addition of trypsin. Overnight digestion at 30°C was performed. Peptides were eluted from gel pieces twice using 100 µl of 1.67% (v/v) TFA in acetonitrile, incubating at 37°C for 15 min with mixing (1200 rpm) each time. Supernatants were combined in 1.5 ml low-binding Eppendorf tubes, frozen on dry ice and vacuum dried.

### Enrichment of crosslinked peptides with MCX SPE cartridges

One of the tryptic digested samples from each experimental set (Rab-8A, PPM1H, and mixture of Rab-8A and PPM1H) were re-dissolved in 1 mL aqueous solution containing 4% (v/v) H_3_PO_4_. Samples were then sonicated in water bath for 30 minutes. MCX cartridge was used to enrich the crosslinked peptides according to the previous protocol[27]. The MCX cartridges were first washed with 2 mL MeOH, then re-conditioned with 2 mL aqueous solution containing 4% (v/v) H_3_PO_4_ prior to sample loading. Samples were then individually loaded onto the MCX cartridges, followed by washing with 500 µL aqueous solution containing 4% (v/v) H_3_PO_4_ and 500 µL of 10% (v/v) MeOH solution with 0.1% (v/v) FA. Low-charged peptides were removed by 500 µL solution composed of 500 mM NH_4_OAc in 40% (v/v) MeOH solution with 0.1% (v/v) FA. High-charged peptides were eluted by 700 µL solution containing 2000 mM NH_4_OAc in 80% (v/v) MeOH solution with 0.1% (v/v) FA. The eluted samples were further dried by SpeedVac.

### LC MS/MS analysis of crosslinked peptides

All dried samples were re-suspended in 30 µL solution containing 3% (v/v) ACN and 0.1% FA, and further sonicated in water bath for 30 minutes. Liquid chromatography tandem mass spectrometry (LC MS/MS) experiment was performed on an Ultrimate 3000 RSLC nano-HPLC system coupled to an Orbitrap Exploris^TM^ 480 mass spectrometer. 3 – 14 µL solution from each sample was loaded onto the nano-HPLC system individually. Peptides were trapped by a precolumn (Acclaim PepMap^TM^ 100, C18, 100 µm x 2 cm, 5 µm, 100 Å) using aqueous solution containing 0.1% (v/v) TFA. Peptides were then separated by an analytical column (PepMap^TM^ RSLC C18, 75 µm x 50 cm, 2 µm, 100 Å) at 45°C using a linear gradient of 1 to 35% solvent B (solution containing 80% ACN and 0.1% FA) for 90 minutes, 35 to 85% solvent B for 5 minutes, 85% solvent B for 10 minutes, 85% to 1% solvent B for 1 minute, and 1% solvent B for 14 minutes. The flow rate was set at 300 nL/min for all experiments. Data were acquired with data-dependent MS/MS mode. For each MS scan, the scan range was set between 375 and 1500 *m/z* with the resolution at 120,000 and 300% automatic gain control (AGC) was used. The maximum injection time for each MS scan was 100 ms. The 10 highest abundant peptides with charge state between 2 and 8 as well as intensity threshold higher than 1.0e+4 were then isolated with a 1.2 Da isolation window sequentially. Stepped HCD with normalized collision energy of 27, 30, and 33% was applied to fragment the isolated peptides. For each MS/MS scan, the resolution was set at 15,000 with a normalized AGC at 200%. The maximum injection time was set at 250 ms. Dynamic exclusion with 60 s duration and 2 ppm window was enabled for the experiment.

### Identification of crosslinked PPM1H/pRab8a peptides

The .RAW files obtained from the LC MS/MS experiments were converted into .mgf files using RawConverter software[42]. The .mgf files were submitted to search using MeroX software against PPM1H and Rab8a protein sequences to identify potential crosslinked peptides[42, 43]. Digestive enzyme, trypsin, trypsin and AspN, or trypsin and GluC, were selected according to the experimental setup. 3 maximum missed cleavages with peptide length ranged from 3 to 50 were applied. Carbamidomethylation at cysteine residues was set as fixed modification, while oxidation at methionine residue and deamidation at asparagine residue were included in variable modification. DSBU crosslinker was selected with specificity crosslinked sites at lysine, serine, threonine, and tyrosine residues. 10 ppm and 20 ppm were used to filter the mass error in precursor ion (MS1) and fragment ion (MS2) scans. Only ions with signal-to-noise ratio high than 2 were used for database search. RISEUP searching mode was applied, minimum 2 fragments per peptide and 5% false discovery rate (FDR) were required for a crosslinked peptide identification. Potential crosslinked peptides with score higher than 50 were then manually check to guarantee the cleavage information obtained from the MS experiment could identify only a single crosslinked site in a peptide. A detailed protocol describing the crosslinking and mass spectrometry analysis methodology has been reported (dx.doi.org/10.17504/protocols.io.bv2en8be).

### Mass photometry analyses of PPM1H

Mass photometry was performed using OneMP mass photometer (Refeyn Ltd, Oxford, UK) and data acquisition performed using AcquireMP (Refeyn Ltd, Oxford, UK). Microscope coverslips (No. 1.5, 24 × 50, VWR) and Grace Bio-Labs reusable CultureWellTM gaskets (3 mm x 1 mm, well capacity 3-10 ul, Sigma-Aldrich, GBL103250-10EA) were cleaned 3 times sequentially in 100% isopropanol and Milli-Q H_2_O, followed by drying with a stream of compressed air. The gasket was carefully placed and pressed onto the centre of the cover slip to assemble the chamber. Prior to mass photometry measurements, protein marker (NativeMark Unstained protein standard, LC0725, ThermoFisher) was diluted 50x in the working buffer (40 mM HEPES pH 7.5). To find focus, 15 µl of buffer was first pipetted into the well, and the focal position was identified as described in the manual. 2.5 µl of protein marker was pipetted into the same well containing the working buffer. A movie of 10000 frames was recorded and distribution peaks of the protein standards were fitted with Gaussian functions to obtain the average molecular mass of each distribution component and used to generate the standard curve. Protein stocks were diluted in working buffer to 40 nM final concentration in final volume of 15 µl. For each acquisition the diluted protein was pipetted into a new well and movies of 10000 frames were recorded. Each sample was measured least three times. Images were analyzed using DiscoverMP (Refeyn Ltd, Oxford, UK).

## Acknowledgements

We thank the excellent technical support of the MRC-Protein Phosphorylation and Ubiquitylation Unit (PPU) DNA Sequencing Service (coordinated by Gary Hunter), the MRC-PPU tissue culture team (coordinated by Edwin Allen), MRC PPU Reagents and Services antibody and protein purification teams (coordinated by Hilary McLauchlan and James Hastie), MRC-PPU mass spectrometry facility (coordinated by Dr. Renata F. Soares) for help in the maintenance of the LC and mass spectrometers. This work is based upon research conducted at the Northeastern Collaborative Access Team beamlines, which are funded by the National Institute of General Medical Sciences from the National Institutes of Health (P30 GM124165). The Eiger 16M detector on the 24-ID-E beam line is funded by a NIH-ORIP HEI grant (S10OD021527). The resources of the Advanced Photon Source, a U.S. Department of Energy (DOE) Office of Science User Facility, is operated for the DOE Office of Science by Argonne National Laboratory under Contract No. DE-AC02-06CH11357. This research also used the FMX beamline of the National Synchrotron Light Source II, a US Department of Energy (DOE) Office of Science User Facility operated for the DOE by Brookhaven National Laboratory under Contract No. DE-SC0012704. We thank Suzanne Pfeffer for helpful advice and discussions.

## Intellectual property rights notice

“This research was funded in part by Aligning Science Across Parkinson’s [Grant ID: ASAP-000463] through the Michael J. Fox Foundation for Parkinson’s Research (MJFF). For the purpose of open access, the authors have applied a CC-BY public copyright license to the author accepted manuscript version arising from this submission.”

## Author Contributions

D.R.A. and A.R.K. designed experiments and supervised the work. D.W., A.R.K., A.K. and K.B. produced recombinant proteins. D.W. and A.R.K. performed crystallization and A.R.K. determined the structures. K.B. designed and carried out cellular and *in vitro* kinetic assays. P.L. and Y.P.Y.L. performed the crosslinking study. D.W., K.B., P.L., Y.P.Y.L., D.R.A and A.R.K. wrote the manuscript.

## Data Availability

The structures of PPM1H phosphatase have been deposited in the Protein Data Bank with codes 7kpr, 7l4j, 7l4i, and 7n0z. All primary data associated with each figure has been deposited in the Zenodo data repository with a doi number dx.doi.org/10.5281/zenodo.5045023. The mass spectrometry proteomics data have been deposited to the ProteomeXchange Consortium via the PRIDE partner repository with the dataset identifier PXD026367.

## Expanded View Figure Legends

**Figure EV1:**
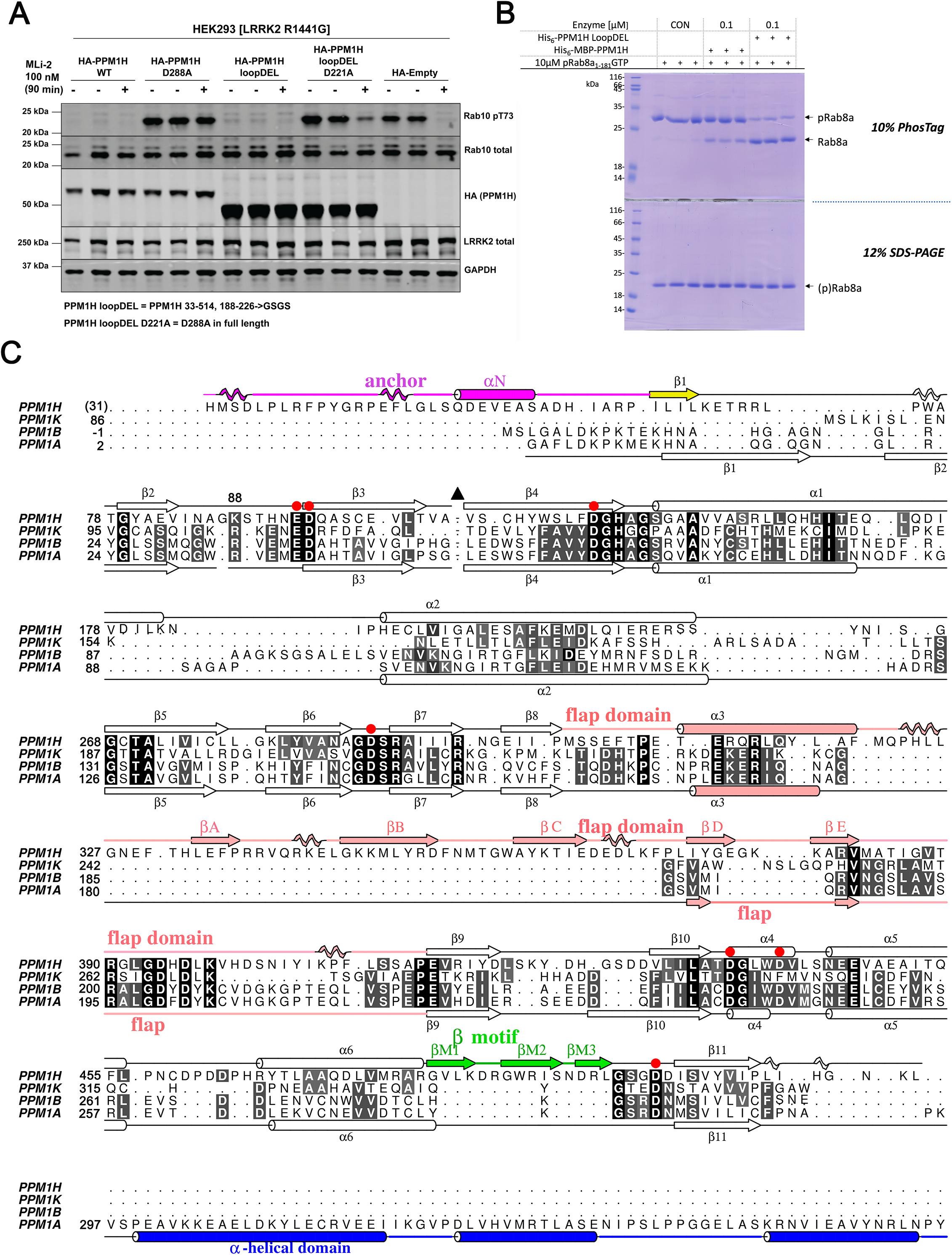
The ‘loop deletion’ variant of PPM1H is active in vitro and in cells. **(A)** HEK293 cells overexpressing indicated constructs were treated ±200 nM MLi-2 for 90 min and then lysed. 10 µg whole cell lysate was subjected to immunoblot analysis with the indicated antibodies at 1 µg/ml final concentration and membranes were analysed using the OdysseyClx Western Blot imaging system. Each lane represents cell extract obtained from a different dish of cells (two replicates per condition without MLi-2 treatment, one replicate per condition with MLi-2 treatment). The ‘loopDEL’ variant is the segment 33-514 with the region 188-226 replaced by the sequence ‘GSGS’. **(B)** *In vitro* assay of PPM1H activity using a PhosTag gel. Substrate pRab8a (GTP form, 10μg) was incubated with WT PPM1H ±loopDEL for 15min at room temperature. The full-length variant was fused to maltose-binding protein (MBP). The ‘loopDEL’ variant was from 33-514 and was used for crystallization studies. A conventional 12% SDS-PAGE gel is shown in parallel lanes below the PhosTag gel. **(C)** Structure-based sequence alignment of human PPMs from their associated PDB files using Chimera software[44]. The PDB codes are 4ra2 (PPM1A; [45]), 2p8e (PPM1B; [46]), and 2iq1 (PPM1K; [46]). Residue His31 of PPM1H is in brackets since the first two residues (His-Met) arise from a cloning artifact. Secondary structures of PPM1H and PPM1A are above and below the sequences, respectively. Colours of secondary structures correspond to the scheme in Fig 1. The black triangle is a loop in PPM1H (188-226) that has been removed to simplify the alignment. The C-terminal α-helical domain of PPM1A is blue. Red circles are aspartate residues that directly coordinate metal ions.

**Figure EV2:**
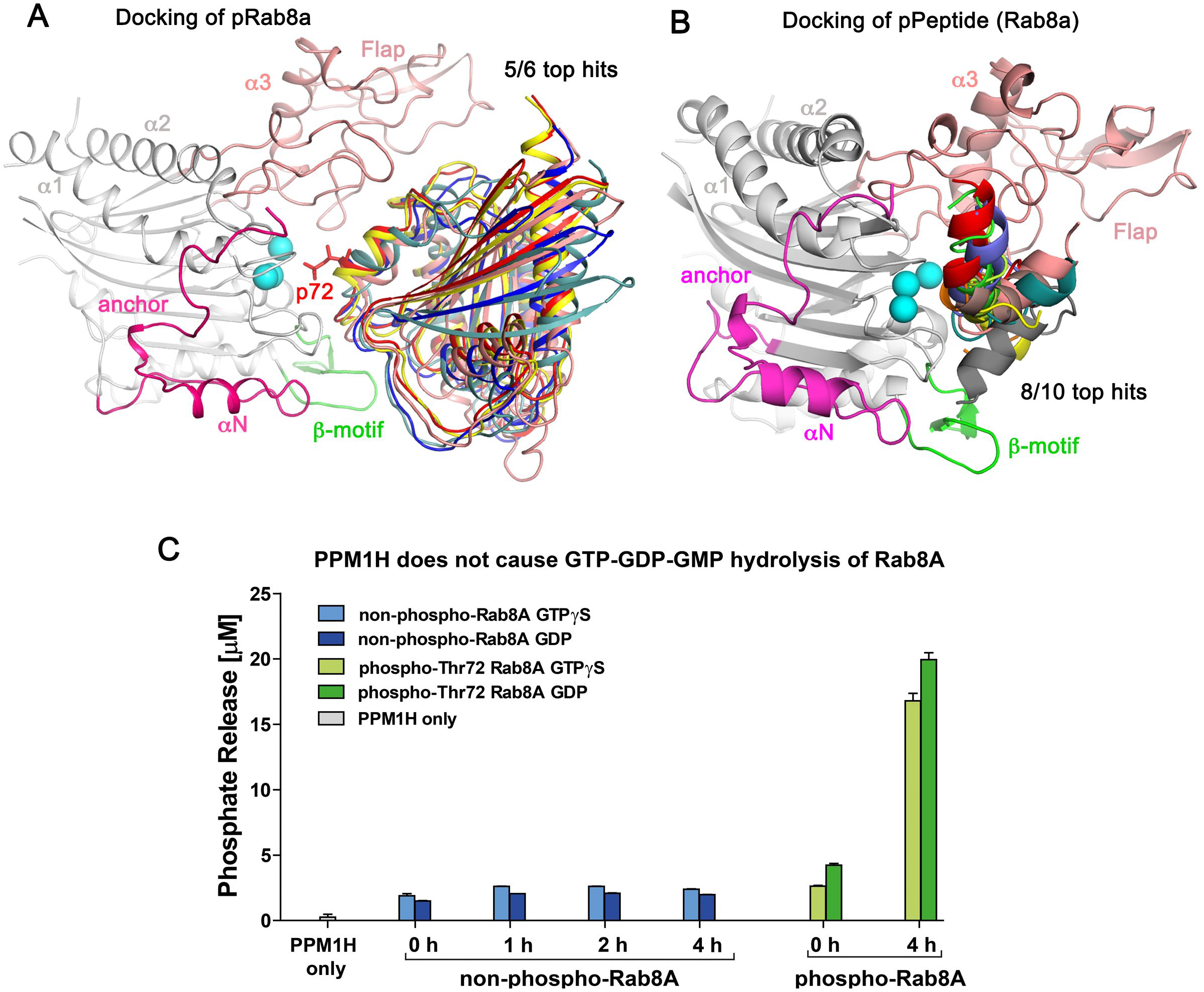
**(A)** Docking of 5/6 top solutions using Haddock software. A stick model for pThr72 (p72, red) is shown for one of the hits, adjacent to the Mn^2+^ ions. All of these docking poses pack against the flap domain. **(B)** Docking of the switch 2 phosphopeptide against MnPPM1H^WT^-LD. The solutions are in addition to those shown in Fig 2 and are shown for completeness of the docking results. However, these 8 poses are sterically incompatible with the active site in the context of the full-length pRab8a protein. **(C)** 50 nM recombinant wild-type PPM1H was incubated with 32 µM of non-phospho- and pThr72 phosphorylated Rab8a (GTPγS or GDP) for 1, 2 or 4 h. Phosphate release was measured as described in Materials and Methods.

**Figure EV4:**
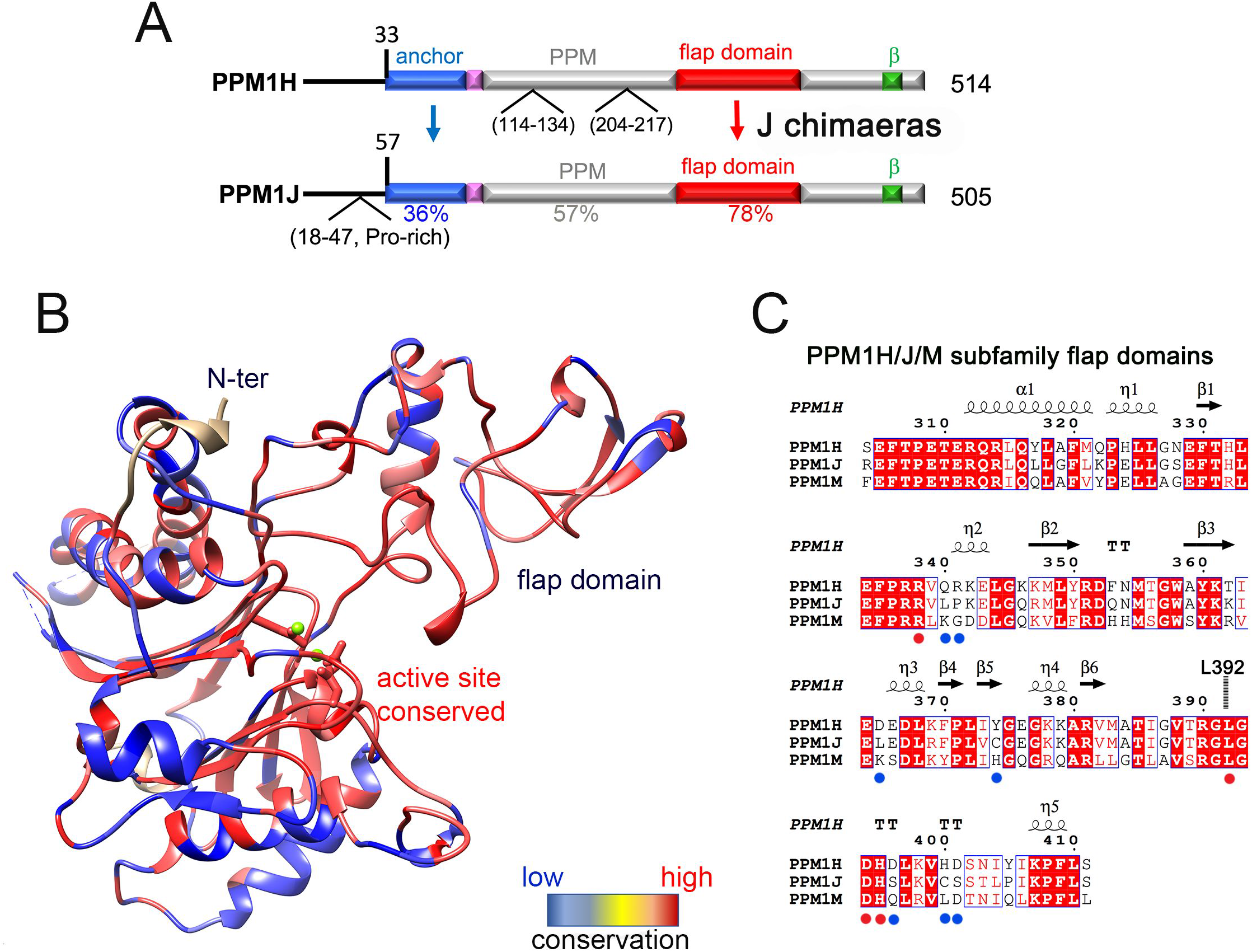
Design of PPM1H/J chimeras to probe substrate specificity. **(A)** Domain organization of PPM1H and PPM1J with locations of predicted flexible loops. Sequence identities within domains are indicated below the alignment. **(B)** Heat map of PPM1H and PPM1J sequence diversity superimposed onto the ribbon model of PPM1H. The rectangular bar displays the variation from low (blue) to high (red) sequence identities. **(C)** Sequence alignment of the flap domains of PPM1H and PPM1J. Secondary structure annotations of PPM1H are above the alignment. Blue and red circles below the alignment indicate conserved and non-conserved residues subjected to site-directed mutagenesis.

